# Cellular NAD^+^ availability and redox state constrain developmental speed in the *Drosophila* eye

**DOI:** 10.1101/2025.09.02.673715

**Authors:** Nisha Veits, Yuting Guo, Jingjing He, Khalil Mazouni, Ivan Nemazanyy, Martin Bres, Cara Picciotto, Claire Mestdagh, Yan Yan, Francois Schweisguth

## Abstract

The cellular and biochemical processes that limit the speed at which embryos develop, tissues form, and cells differentiate remain largely unknown. Using the speed of progression of a differentiation front in the developing eye of *Drosophila* as a proxy for developmental speed, we identified genetic perturbations that slowed down the progression of this front. Inhibiting the Electron Transport Chain (ETC), and more generally energy production in mitochondria, resulted in reduced developmental speed. Defective ETC activity led to increased NADH/NAD^+^ ratio whereas ATP levels remained constant due to a compensatory increase in glycolysis. Using targeted perturbations, we found that the metabolic state of the cells ahead of and/or at the moving front of differentiation determined its speed. Genetic and diet-based perturbations of the NAD^+^ metabolism pathway indicated that developmental speed was limited by NAD^+^ availability in these cells. Thus, developmental speed appeared to be constrained by the cellular redox and the demand for NAD^+^ in *Drosophila*.

## Introduction

Time is an inherent property of living systems, and many biological processes have defined time scales. For instance, the one-cell embryo of *Drosophila* produces a complex larva within a single day at 25°C. However, while it is commonly observed that embryos and tissues develop at a reproducible tempo for defined environmental conditions, little is known about what sets the speed at which ordered, interacting and non-reversible steps occur during development (Ebisuya and Briscoe, 2018). At the organismal level, developmental speed can be influenced by nutrition and temperature (Gillooly et al., 2002), indicating that metabolism constrains and influences the speed of development (Ghosh et al., 2023). Indeed, differences in mitochondrial activity, energy metabolism, protein stability and/or gene expression were recently shown to underly the species-specific differences in developmental speed between different mammalian embryos (Diaz-Cuadros et al., 2023; Iwata et al., 2023; Lázaro et al., 2023; Matsuda et al., 2020; Matsuda et al., 2025; Nakanoh et al., 2024; Porte et al., 2024; Rayon et al., 2020). Notably, the inhibition of the mitochondrial Electron Transport Chain (ETC) was shown to reduce the speed of the segmentation clock in an *ex vivo* stem cell-based assay mimicking somitogenesis in the mouse (Diaz-Cuadros et al., 2023). The ETC uses the energy produced by the transfer of electrons from donors, e.g. reduced nicotinamide dinucleotide (NADH), to acceptors to pump protons in the intermembrane space of mitochondria, thereby generating a proton gradient that drives the production of Adenosine TriPhosphaste (ATP). Since the first protein complex of the ETC, complex I (cI) oxidizes NADH, inhibiting its activity increases the NADH/NAD^+^ ratio, affecting in turn the activity of many enzymes regulating catabolism and anabolism. Since various perturbations changing the NADH/NAD^+^ ratio altered the rate of somitogenesis in this *ex vivo* assay, developmental speed was proposed to be regulated by the NADH/NAD^+^ ratio in this context (Diaz-Cuadros et al., 2023). Beyond cellular redox state, other processes have been proposed to modulate developmental speed in mammalian embryos (Diaz-Cuadros et al., 2023; Iwata et al., 2023; Lázaro et al., 2023; Matsuda et al., 2020; Matsuda et al., 2025; Nakanoh et al., 2024; Porte et al., 2024; Rayon et al., 2020), raising the possibility that distinct molecular processes contribute to various extents in the regulation of developmental speed in different tissues and/or species (Matsuda et al., 2025).

The *Drosophila* eye is an attractive model system to study developmental speed. The adult fly eye comprises ~750 light-receiving multicellular units, or ommatidia, that are arranged in a crystal-like array. It develops from an eye primordium in the eye-antenna imaginal disc (Ready et al., 1976). This eye primordium differentiates in a progressive manner as a moving differentiation front, forming a morphogenetic furrow (MF), sweeps through the disc epithelium from posterior to anterior over a 2.5 day-period to produce ~30 rows of regularly spaced ommatidia (Roignant and Treisman, 2009) (Fig 1A-B’’’). Cells located anterior to the front remain undifferentiated and proliferate whereas cells posterior to the furrow differentiate and form ommatidia. This front moves at a relatively constant speed of ~0.6 rows per hour (hr) (Spratford and Kumar, 2015; Tsao et al., 2016), or ~3 μm/hr (Wartlick et al., 2014), with a new row of ommatidia produced at each pulse of Atonal, the proneural transcription factor that controls eye differentiation (Couturier et al., 2024). A self-propagating mechanism involving signals produced by differentiating cells underlies the progression of the front (Couturier et al., 2024; Greenwood and Struhl, 1999; Ma et al., 1993; Roignant and Treisman, 2009). Thus, the progression speed of the differentiation front can be used as a proxy to study the speed of developmental patterning. This system offers key experimental advantages. First, the temporal progression of patterning and differentiation can be easily mapped in space. It can therefore be studied in fixed tissues. Second, the molecular mechanisms underlying the progression of the front is well understood. Third, genetic tools allow for targeted perturbations of developmental speed in the eye. Indeed, compartment-specific perturbations make it possible to study relative changes in developmental speed within a single eye disc, thereby overcoming the systemic influence of extrinsic factors, such as temperature, nutrition, feeding behavior, infection and other stresses on developmental speed. In this approach, the unperturbed compartment of the disc can be used as an internal control for any physiological change acting at the organismal level. Thus, the fly eye appears as a good model system to study developmental speed at the tissue level in the context of the living organism.

**Figure 1:**
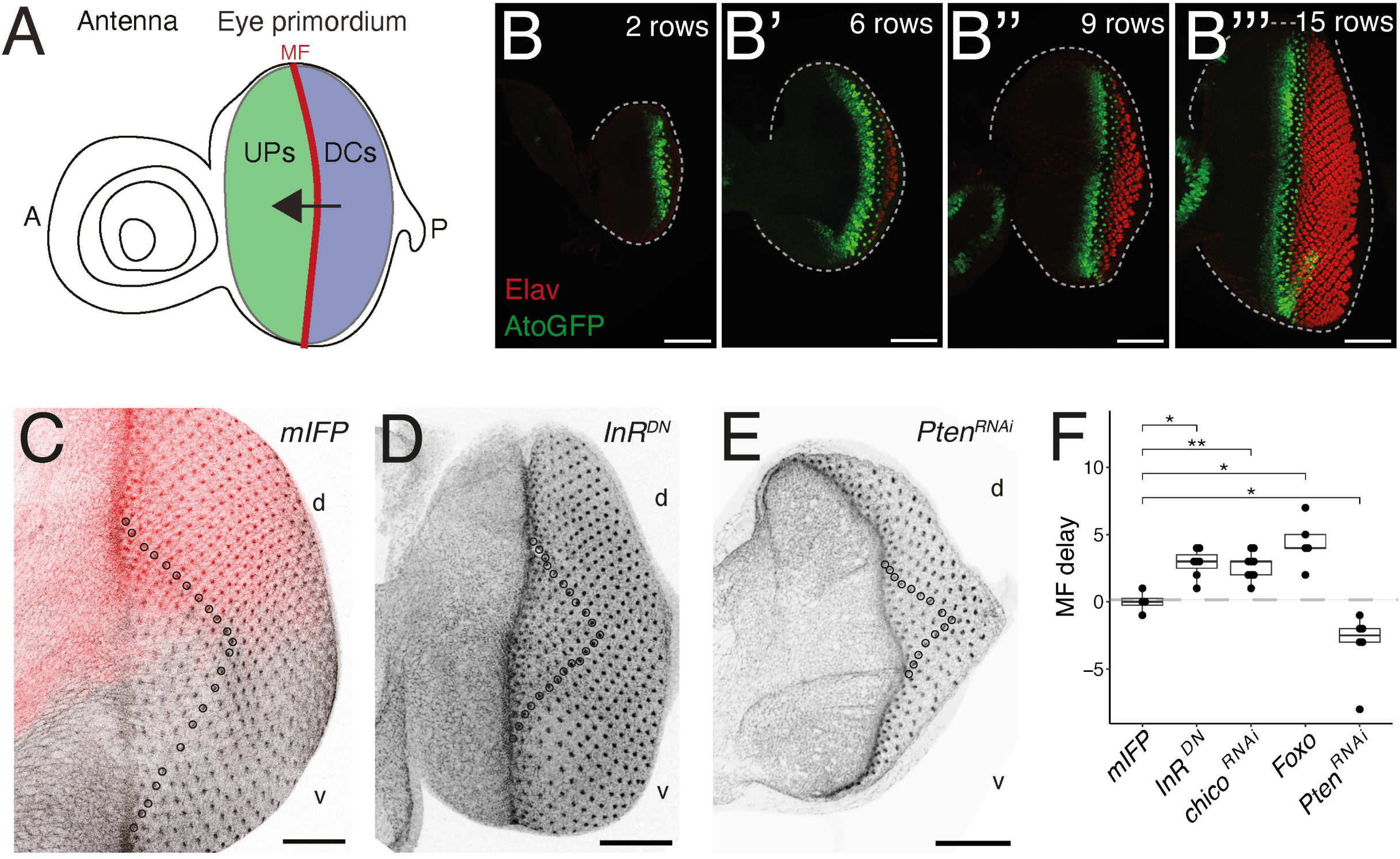
A simple assay for front progression. **A)** Schematic of an eye-antenna imaginal disc. The MF (red) travels through the eye primordium from posterior (P) to anterior (A; arrow) as Undifferentiated Progenitors (UPs; green) become Differentiated Cells (DCs; blue). **B-B’’’)** Time series of eye discs stained for Atonal (AtoGFP, green) marking the MF and Elav (red) marking DCs. The number of ommatidial rows was used as a proxy of developmental time. Note that disc growth correlates with MF progression. **C,D,E)** Mapping time in space: ommatidia identified using Ecad (black) were counted (circles) from the midline to the MF in dorsal (*d*) and ventral (*v*) compartments in *mirr>mIFP* (C; mIFP, red, marked *d* cells), *mirr>InR^DN^*(D) and *mirr>Pten^RNAi^*(E) eye discs. **F)** Relative changes in MF progression, measured as the difference in row numbers in the *v* and *d* compartments, upon expression of InR^DN^ (2.9 +/−1.0, n=6), *chico^RNAi^*(4.4 +/−1.8, n=7), Foxo (2.7 +/−1.0, n=5) and *Pten^RNAi^*(−3.2 +/−2.8, n=6). Expression of mIFP (0.0 +/−0.8, n=4) served as a negative control. While InR^DN^, *chico^RNAi^* and Foxo led to a MF delay, *Pten*^RNAi^ resulted in increased MF progression. Wilcoxon tests: * < 0.05; ** < 0.01. Scale bars, 50µm.

Here, we used a simple *in vivo* assay to identify RNAi-based perturbations that affected the timely progression of the MF. This approach showed that the mitochondrial ETC and Oxidative Phosphorylation (OxPhos) activities were required for the proper progression of the differentiation front. Using fluorescent metabolic sensors and metabolomic approaches, we found that the loss of ETC activity increased the NADH/ NAD^+^ ratio but did not detectably change the level of ATP. ATP levels were maintained by increased glycolysis upon loss of ETC activity which was associated with increased *Lactate dehydrogenase* (*Ldh*) gene expression, likely to promote NAD^+^ regeneration. Indeed, expression of an heterologous NADH oxidase and addition of NAD^+^ precursors in the diet indicated that the rate of NAD^+^ regeneration from NADH appeared to limit the rate of progression of the MF upon loss of ETC activity. Moreover, the biosynthesis of NAD^+^ by the salvage pathway was found to be important for proper developmental speed in the eye, and while changes in developmental speed often correlated with changes in NADH/ NAD^+^ ratio, exceptions were noted in cells for which energy metabolism played a key role for the progression speed of the front. Our data therefore indicated that the demand for NAD^+^ constrained developmental speed in the developing fly eye.

## Results

### An assay for developmental speed in the eye

The front of differentiation moves at a relatively constant rate, with a new row of ommatidia forming every ~100min at 25°C (Spratford and Kumar, 2015; Tsao et al., 2016) (Fig 1A-B’’’). What determines this speed is not known. Since the speed of development at the organismal level varies with systemic variations in circulating nutrients, hormones and other secreted factors, the speed of progression of the MF should vary with nutrition, stress, social interactions and other environmental factors (Cassidy et al., 2019). This variability therefore complicates the analysis of the progression speed of the MF at the population level. To overcome this issue, we studied here relative differences in speed within individual eye discs by measuring the relative progression of the MF in experimental dorsal (*d*) cells expressing the *mirror* (*mirr*) *Gal4* driver relative to control ventral (*v*) cells. A difference in progression speed was inferred by counting the number of ommatidial rows produced over a given time in the *v* and *d* compartments of the same disc (Fig 1C). Scoring fewer rows of ommatidia in *d* cells relative to *v* cells would indicate that the progression of the MF is delayed in the *d* compartment, whereas scoring more rows would indicate an increase in speed. Thus, this assay does not measure absolute speed and may rather reveal relative differences in speed. In addition, this simple assay should be faster than population-based assays used earlier (Spratford and Kumar, 2015; Tsao et al., 2016; Wartlick et al., 2014), therefore allowing us to test many genetic perturbations.

To validate this assay, we tested the Insulin signaling pathway that is known to regulate cell growth, cell proliferation and differentiation timing in the *Drosophila* eye (McNeill et al., 2008, Cassidy et al., 2019). Using a dominant-negative (DN) form of the Insulin receptor (InR), RNAi-mediated silencing of a conserved and essential InR substrate (Chico), as well as the over-expression of Foxo, a transcription factor inhibited by InR signaling, we found that lowering Insulin signaling activity delayed the progression of the MF (Fig 1D,F and Table S1). In contrast, increasing InR signaling by silencing the Phosphatase and TENsin homolog (Pten) led to increased progression of the MF (Fig 1E,F and Table S1). These results indicated that developmental speed in the eye positively correlates with InR signaling, hence validating our simple assay. Of note, a delay in MF progression in larvae did not produce an easily scorable phenotype in the adult eye (Fig S1A,B), implying that developmental speed defects cannot be easily screened in adult eyes. In summary, since scoring a small number of eye discs was sufficient to detect a statistically significant difference in MF progression, this simple assay should in principle allow us to identify perturbations that reduce or increase the rate of development.

### Delayed progression of the MF upon energy metabolism perturbations

We then used this assay to look for RNAi-based perturbations associated with altered developmental speed. To probe various aspects of cellular metabolism, including energy metabolism, stress responses and biosynthesis and turnover of macromolecules, we selected and tested 166 genes (Table S1). This screen identified several genes encoding subunits of the complexes cI, cIV and cV of the ETC (Fig 2A-C), including *ND-42*, a cI accessory subunit known as NDUFA10 in mammals (Kampjut and Sazanov, 2022; Wirth et al., 2016) and *bellwether* (*blw*), the alpha subunit of the mitochondrial F_0_F_1_-ATP synthase complex (cV). The MF delay seen upon *ND-42^RNAi^* was suppressed by the expression of the yeast alternative NADH oxidase *NDI1* which can bypass a loss of cI activity (Seo et al., 1998) (Fig 2D). This showed that the *ND-42^RNAi^* MF delay phenotype resulted from a loss of cI activity, implying that loss of OxPhos activity delayed the progression of the MF (Fig 2C). Additionally, the RNAi-mediated silencing of the *Carnitine PalmitoylTransferase 2* (*CPT2*) and *pyruvate dehydrogenase* (*pdhb*) genes similarly delayed the progression of the MF (Table S1), suggesting that fatty acid β-oxidation and production of Acetyl-CoA downstream of glycolysis were also important for the timely progression of the MF. In contrast, RNAi-mediated inhibition of many glycolytic and TCA enzymes showed little effect (Table S1). Beyond energy metabolism, a few enzymes involved in other metabolic pathways scored positively in our assay (Table S1). However, no perturbation associated with increased furrow progression was identified (beyond *Pten^RNAi^*, Fig. 1E). In sum, our results strongly indicated that energy production by mitochondria is key for the timely progression of the MF in *Drosophila*.

**Figure 2:**
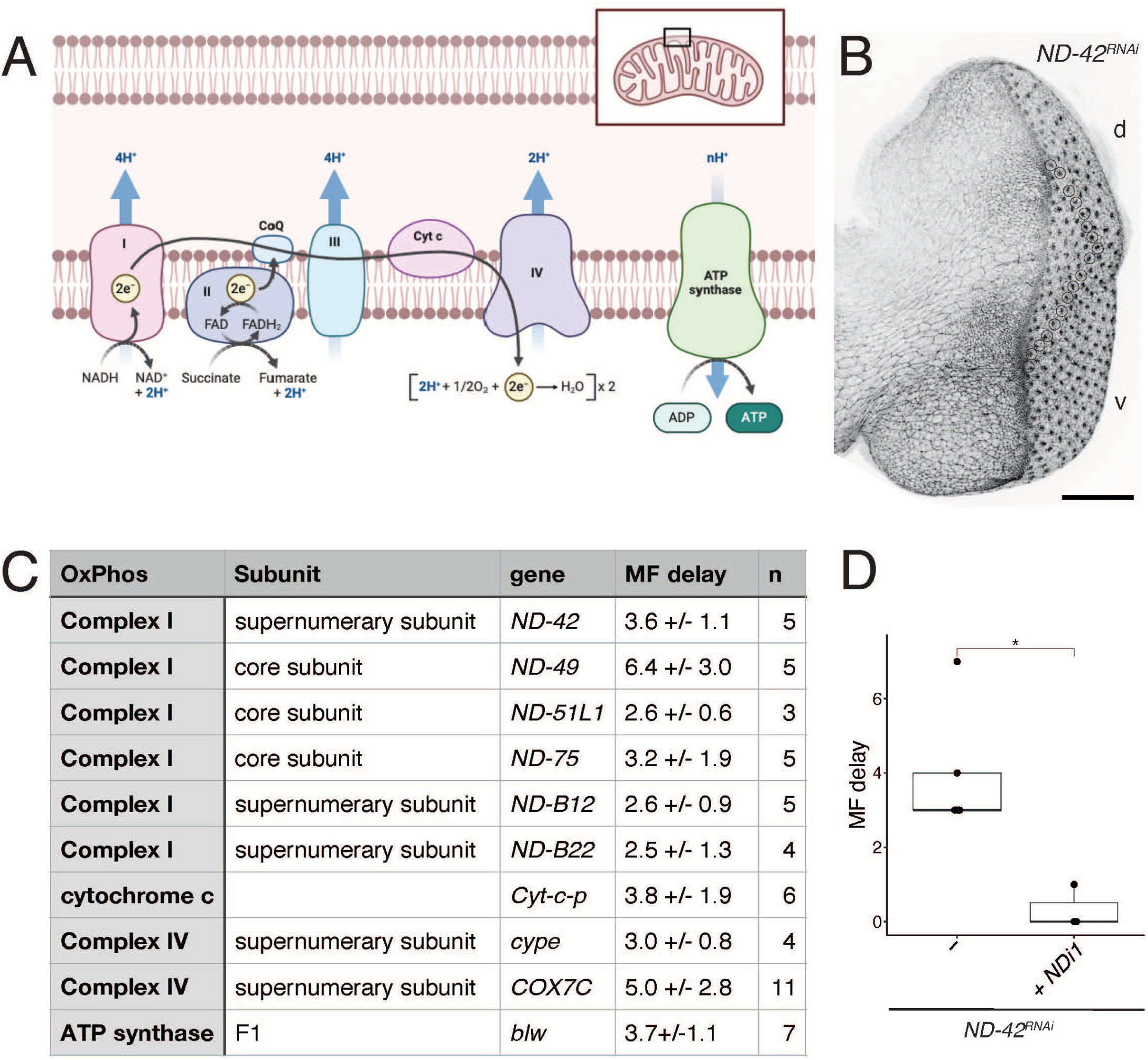
Delayed progression of the front upon ETC inhibition. **A)** Schematic of the ETC (adapted from BioRender.com). **B)** A *mirr>ND-42^RNAi^* eye discs (Ecad, black) showing a difference in the number of ommatidial rows (circles), hence a delay in MF progression. **C)** Results of the MF delay assay for selected ETC genes. **D)** Expression of the yeast alternative NADH oxidase *NDI1* suppressed the *ND-42^RNAi^* MF delay phenotype. Wilcoxon test:* < 0.05. Scale bar, 50µm.

### Reduced speed of front progression upon cI inhibition

To further test whether the delayed progression of the MF resulted from reduced speed, we designed a recombination-based assay that allowed us to infer the speed of the differentiation front from fixed eye discs. In brief, a heat-induced recombination-based RFP-to-GFP switch followed by a chase period allowed us to score the number of GFP-positive rows produced in a defined amount of time (Fig 3A,B). This assay was validated by showing that speed varied with temperature (Fig S1C). We next measured the average speed of the front in control larvae (Fig 3C) and found that the MF moved at a speed of ~0.6 row/hr, consistent with earlier reports (Spratford and Kumar, 2015; Tsao et al., 2016). Analysis of *mirr>ND-42^RNAi^* discs showed that the silencing of the *ND-42* gene led to a ~30% reduction in relative speed (0.25 vs 0.39 row/hr in Fig 3D). We therefore propose that the observed delay in MF progression likely resulted from a slow speed of the moving front of differentiation. Of note, the speed measured in control *v* cells of *mirr>ND-42^RNAi^* discs (0.39 row/hr) was lower than the speed measured in control larvae (~0.6 row/hr), suggestive of a developmental delay in *mirr>ND-42^RNAi^* larvae. We next addressed whether cI inhibition perturbed eye patterning and differentiation by studying the pattern of expression of key regulators of eye differentiation in *mirr>ND-42^RNAi^* discs. We found that eye patterning remained largely unchanged upon cI inhibition (Fig S1C-D’). We therefore conclude that cI inhibition slowed the progression of the MF.

**Figure 3:**
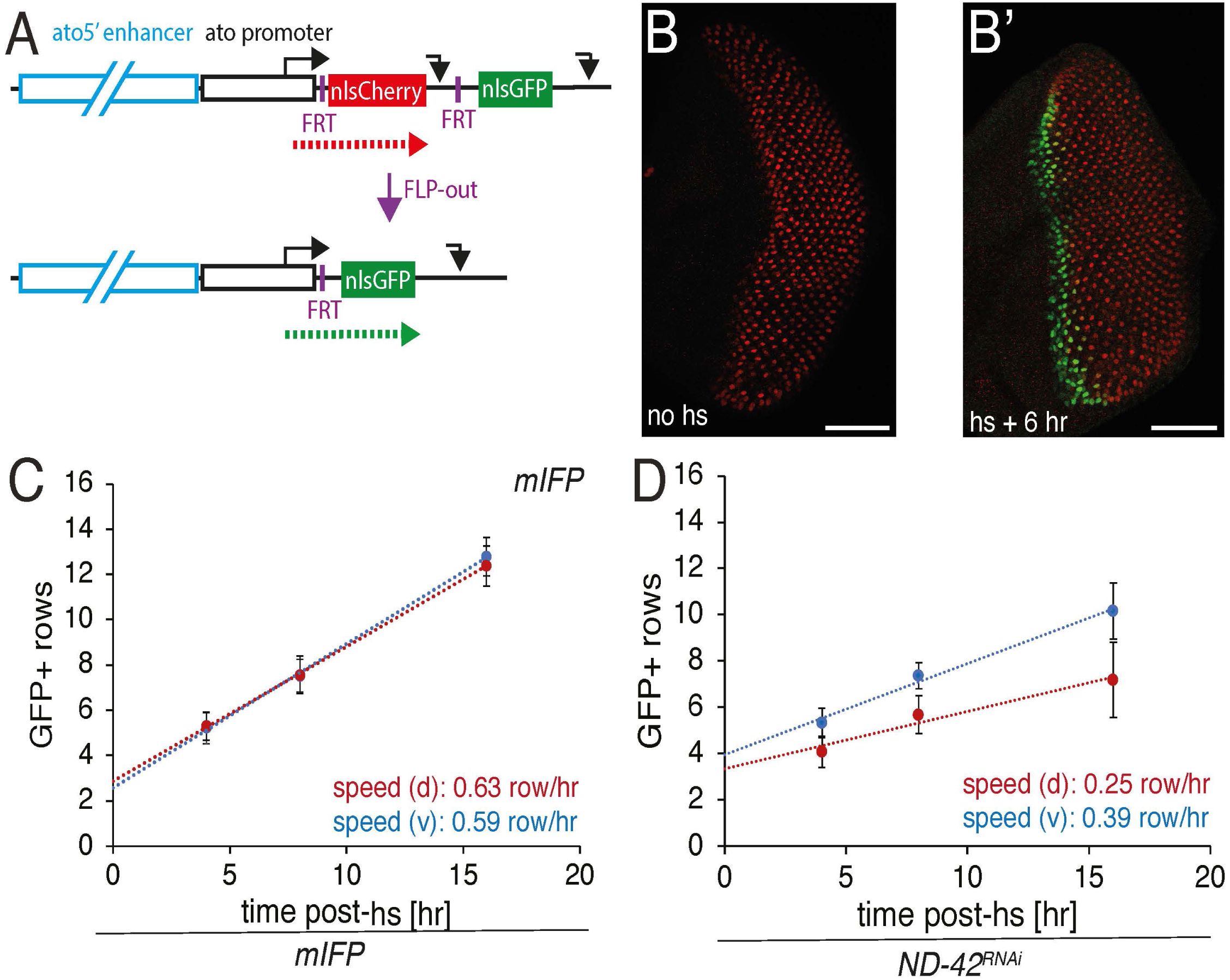
Reduced speed of progression of the front upon ETC inhibition. **A-B’)** Recombination-based strategy to infer MF speed from fixed samples (A). The Flp-mediated excision of nuclear mCherry upon heat-shock (hs) triggers a mCherry-to-GFP switch in gene expression. The *ato5’* enhancer directs the expression of mCherry (or GFP) at the front. Prior to recombination, mCherry produced at the front was stable enough to mark all ommatia (B). Following recombination, new rows of GFP-positive/mCherry-negative omatidia are produced (B’). The speed of progression of the front was inferred by scoring the number of GFP-positive rows produced in the time interval from hs to fixation. Scale bars, 50µm. **C,D)** The speed of progression of the front was determined as the slope of the regression line produced using the number of GFP-positive rows scored 4, 8 and 16 hr after hs. In control *mirr>mIFP* discs, the measured speed was ~0.6 rows/hr (4 hr, n=44; 8 hr, n=25; 16 hr, n=37). No speed difference was observed between *d* and *v* cells in control discs (C). In *mirr>ND-42^RNAi^* discs (D), the front progressed at a slower speed in *d* cells (0.25 rows/hr) relative to control *v* cells (0.39 rows/hr; 4 hr, n=31; 8h, n=29; 16 hr, n=32).

### NADH accumulation upon cI inhibition

The cI is an oxidoreductase located at the inner membrane of mitochondria which accepts electrons from NADH and transfers them to oxygen, thereby converting the produced free energy to pump protons across the inner membrane (Fig 2A). Loss of cI activity therefore decreases the rate of NADH oxidation, leading to NADH accumulation. To examine whether cI inhibition altered the NADH/NAD^+^ ratio, we used the fluorescence sensor SoNar that consists of a NADH/NAD^+^-binding domain fused to a circulated permuted YFP. The excitation spectra of SoNar vary with the relative binding of NADH *vs* NAD^+^ (Zhao et al., 2015). Flies expressing a mitochondria-targeted SoNar in all cells were generated and used to study the mitochondrial NADH/ NAD^+^ ratio in *ex vivo* cultured eye discs. Similar SoNar signals were observed in *d* and *v* cells in control eye discs (Fig 4A), in both Undifferentiated Progenitors (UPs, anterior to the furrow) and Differentiated Cells (DCs, posterior to the furrow), as shown by the *dv* Fold Change (FC) values which were close to 1 (Fig 4A,C). In contrast, the silencing of the *ND-42* gene led to increased NADH/ NAD^+^ ratio values in both UPs as well as DCs (Fig 4B,C). Thus, as expected, cI inhibition led to elevated NADH/ NAD^+^ ratio in mitochondria.

**Figure 4:**
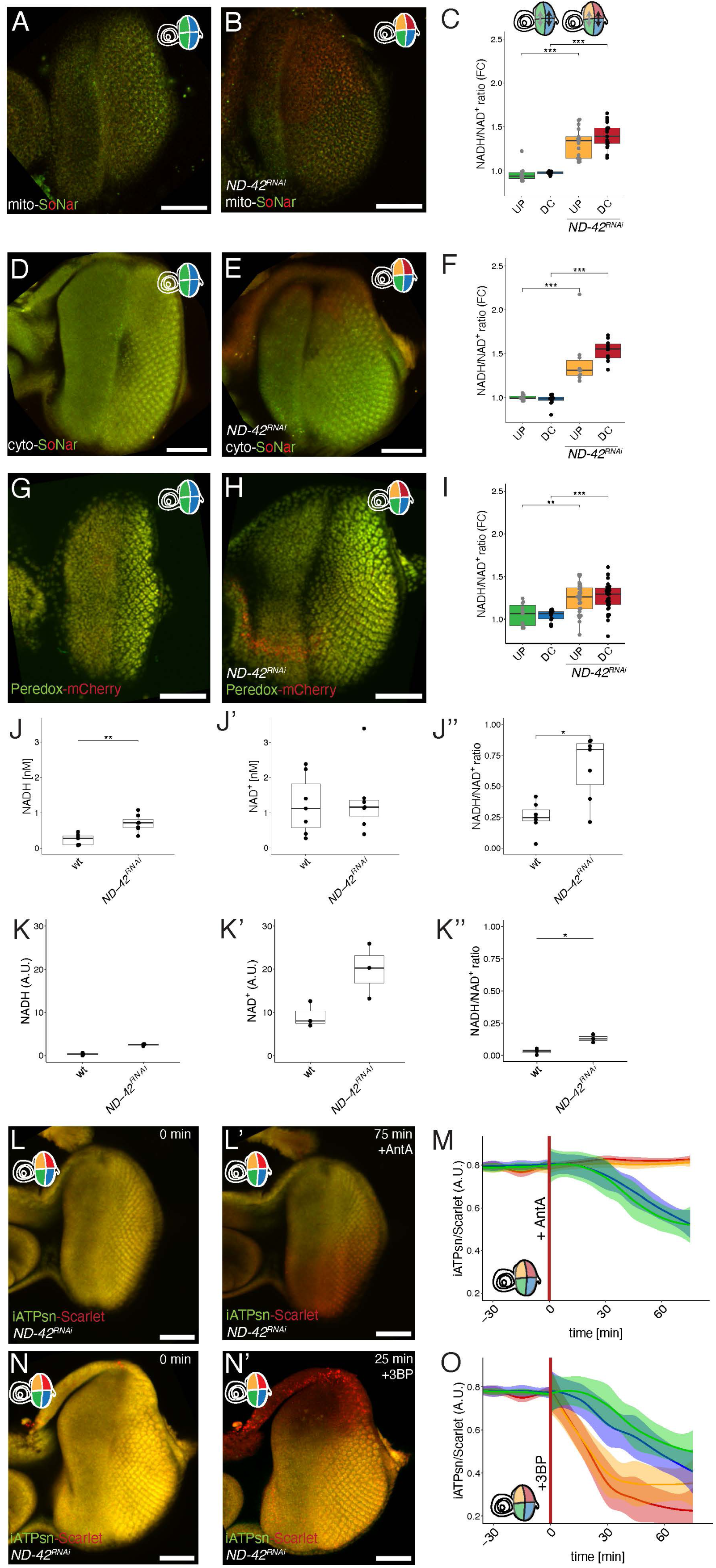
Glycolysis maintained ATP levels constant upon ETC inhibition. **A-C)** Analysis of the mitochondrial NADH/NAD^+^ ratio in control *mirr>mIFP* (A,C) and *mirr>ND-42^RNAi^*+*mIFP* discs (B,C). The *d* cells were identified using the infra-red signal produced by mIFP (not shown). The color code used in C (and throughout the figures) is explained in the schematics shown in A,B (top): control UPs, green; control DCs, blue; *ND-42^RNAi^* UPs, orange; *ND-42^RNAi^* DCs red. The mito-SoNar signals following excitation at 405 and 488nm (in red and green in A,B) were used to measure the NADH/NAD^+^ ratio (C). Since similar ratio values were measured in *d* and *v* cells of control discs, the Fold Change (FC) between ratio values measured in *d* and *v* cells remained close to 1 (UPs: 0.97 +/−0.1; DCs: 0.97 +/−0.06, n=10) In contrast, higher NADH/NAD^+^ ratio values were observed in dorsal UPs and DCs of *ND-42^RNAi^* eye discs (UPs: 1.33 +/−0.16; DCs: 1.4 +/−0.15, n=17), showing that cI inhibition resulted in increased NADH/NAD^+^ ratio in mitochondria. **D-F)** Analysis of the cytosolic NADH/NAD^+^ ratio in control *mirr>mIFP* (D,F) and *mirr>ND-42^RNAi^*+*mIFP* discs (E,F) using cyto-SoNar. The results are shown as in A-C for control (UPs: 0.97 +/−0.06; DCs 0.98 +/−0.07, n=11) and *ND-42^RNAi^* eye discs (UPs: 1.4 +/−0.29; DCs: 1.54 +/− 0.12, n=10). Inhibition of cI led to elevated NADH/NAD^+^ ratio in the cytosol. **G-I)** Analysis of the nuclear NADH/NAD^+^ ratio in control *mirr>mIFP* (D,F) and *mirr>ND-42^RNAi^*+*mIFP* discs (E,F) using a ratiometric Peredox sensor (Peredox, green; mCherry, red). In control eye discs (n=11), similar Peredox/mCherry ratio values were observed in UPs (1.05 +/0.13) and DCs (1.04 +/−0.06). In contrast, silencing the *ND-42* gene led to increased ratio values in both UPs (1.24 +/−0.17) and DCs (1.26 +/−0.17, n=30), indicative of elevated NADH/ NAD^+^ ratio upon cI inhibition. **J-J’’)** Analysis of NADH (J) and NAD^+^ levels (J’), and corresponding NADH/NAD^+^ ratio (J’’), in whole cell lysates from control and *tub^ts^*>*ND-42^RNAi^* eye discs using a NAD^+^/NADH-glo assay. Increased NADH/NAD^+^ ratio largely resulted from increased NADH. **K,K’’)** Quantification of NADH (K), NAD^+^ (K’) and corresponding NADH/NAD^+^ ratio (K’’) in whole cell lysates of control *tub^ts^>+* and *tub^ts^*>*ND-42^RNAi^* eye discs by LC/MS. Increased NADH and NADH/NAD^+^ were observed (the ~2-fold increase in NAD^+^ did not appear statistically significant). **L-O)** Time-course analysis of ATP levels in *mirr>ND-42^RNAi^* eye disc using a ratiometric ATP sensor (iATPsn, green; Scarlet, red in L-N’). Normalized intensity ratios are plotted in M,N. Similar iATPsn/mScarlet ratios were observed prior to drug addition in the *d* and *v* cells of *mirr>ND-42^RNAi^* discs (M, n=12; N, n=9; the mean and standard deviation (s.d.) are shown), as seen also in snapshots taken at t=0 (L,N; ~45 min after the onset of imaging). This indicated that the silencing of *ND-42* did not detectably affect the levels of ATP. However, addition of AntA, a cIII inhibitor, led to a rapid decrease in ATP in ventral UPs and DCs, showing that the ETC was active in these cells (L’,M). In contrast, AntA had no effect on *ND-42^RNAi^* cells, indicating that ATP production did not rely on the ETC in these cells. Addition of 3BP, a glycolysis inhibitor showed that *ND-42^RNAi^* cells, both UPs and DCs, were more sensitive than ventral control cells to glycolysis inhibition (M-O). This showed that the silencing of *ND-42* made UPs and DCs strongly dependent on glycolysis for ATP production. Wilcoxon tests: * < 0.05; ** < 0.01; ***< 0.0001. Scale bars, 50µm.

Since the mitochondrial pools of NAD^+^ and NADH equilibrate with the cytosolic pools, in part via the activity of the Malate-Aspartate shuttle (Hu et al., 2021), we also generated a cytosolic SoNar. Similarly, the silencing of the *ND-42* gene resulted in increased NADH/ NAD^+^ ratio values (Fig 4D-F), showing that cI inhibition changed the NADH/NAD^+^ ratio in both mitochondria and cytosol.

To normalize the measured ratio values by the expression of the sensor, we next used the ratiometric sensor Peredox that consists in a circularly permuted variant of GFP fused to a NAD^+^/NADH-binding protein to monitor the ratio of free NADH over free NAD+ (Hung et al., 2011). Peredox was fused to the red fluorescent protein mCherry that served as an internal control for expression level. Since the cytoplasmic and nuclear pools of NAD^+^ and NADH are readily exchangeable (Cambronne et al., 2016), we used a nuclear version of Peredox-mCherry to report on the redox state of the cell (Diaz-Cuadros et al., 2023). Flies expressing nuclear Peredox-mCherry under the control of a ubiquitous promoter were generated and used to measure changes in the NADH/ NAD^+^ ratio. Analysis of eye discs confirmed that cI inhibition resulted in increased NADH/ NAD^+^ ratio (Fig 4G-I).

We next quantified the levels of NAD^+^ and NADH in whole disc extracts using a NAD^+^/NADH-Glo assay (Fig 4J-J’’) as well as a Liquid Chromatography Mass Spectrometry (LC/MS) approach (Fig 4K-K’’). In both assays, eye-antenna discs expressing the *ND-42^RNAi^* construct in all cells using a *tub-Gal4 tub-Gal80^ts^* driver (Fig S2) were compared with control eye-antenna discs. First, cells from control eye-antenna discs appeared to have 5-10x (Fig 4J,J’; Fig 4K,K’) more NAD^+^ than NADH. Second, elevated NADH/NAD^+^ ratio upon silencing of the *ND-42* gene (2.6x and 4x in Fig 4J’’,K’’) resulted from an increase in NADH levels (3x and 8x in Fig 4J,K) and not from a decrease in NAD^+^. Actually, cI inhibition appeared to result in increased NAD pathway metabolites, including NAD^+^ (2x in Fig 4K’; no change in Fig 4J’; see Table S2), possibly reflecting a compensatory increase in NAD^+^ biosynthesis. Thus, cI inhibition resulted in NADH accumulation and, consistent with earlier results obtained in mammalian cells (Diaz-Cuadros et al., 2023), developmental speed appeared to negatively correlate with the NADH/NAD^+^ ratio in the *Drosophila* eye.

### Glycolysis maintained ATP levels constant upon loss of cI activity

Loss of cI activity should also reduce ATP production in mitochondria. To measure ATP levels, we used a ratiometric fluorescent sensor, iATPsnFR-95A.A119L (Marvin et al., 2024), noted iATPsn hereafter. This sensor is an improved version of the single wavelength iATPsnFR sensor (Lobas et al., 2019). It carries an ATP binding subunit of the F_0_F_1_-ATP synthase complex which was mutated to reduce its affinity to ATP down to the mM range and fused to a circularly permuted sfGFP. To perform ratiometric measurements, this ATP sensor also included the pH-sensitive pHmScarlet, noted Scarlet hereafter. This iATPsn-Scarlet biosensor was expressed ubiquitously in transgenic larvae, and analysis was performed on *ex vivo* cultured eye discs. Analysis of *mirr>ND-42^RNAi^* discs showed no difference in ATP levels between *d* and *v* cells (Fig 4L-O), suggesting that ETC inhibition had no major consequence on the steady-state level of ATP, and that reduced developmental speed did not result from low ATP levels.

We next examined the role of the ETC in ATP production. Inhibiting the activity of the ETC using Antimycin A (AntA), a cIII inhibitor (Xia et al., 1997), led to a rapid decrease in ATP levels in control *v* cells of *mirr>ND-42^RNAi^* discs (Fig 4L’,M). This showed that OxPhos contributes to ATP production in control UPs and DCs cells. In contrast, AntA had little effect on ATP production in *ND-42^RNAi^*-expressing *d* cells (Fig 4L’,M). This showed that the activity of the ETC was already inhibited by the silencing of the *ND-42* gene and that ATP was produced by another metabolic pathway upon loss of cI activity.

Glycolysis, that breakdowns glucose into pyruvate, is another important source of cellular ATP. We therefore hypothesized that glycolysis was up-regulated upon ETC inhibition. To test this, we blocked glycolysis using bromopyruvic acid (3BP), a pyruvate derivate that inhibits two key glycolytic enzymes (Ganapathy-Kanniappan et al., 2009; Li et al., 2022). Inhibition of glycolysis by 3BP led to decreased ATP levels in both control and *ND-42^RNAi^* cells, with a more rapid decrease observed in the *ND-42^RNAi^* cells (Fig 4N-O). This indicated that glycolysis was active in all cells, and that *ND-42^RNAi^*-expressing cells appeared to be more dependent on glycolysis for ATP production. We therefore conclude that increased glycolysis appeared to compensate for the loss of ETC activity, thereby maintaining ATP levels constant.

### Increased glycolysis is a key response to cI inhibition

To further study the cellular response to the loss of ETC activity, we next performed single cell RNA sequencing (scRNAseq). Starting from eye-brain complexes dissected from wild-type and *mirr>ND-42^RNAi^*larvae, we sequenced ~30,000 cells per genotype and identified the eye field as a cluster of ~3,000 *eya*-positive and *sim*-negative cells. Further clustering produced three sub-clusters corresponding to UPs, DCs and peripodial cells based on key marker genes (Fig S3A-B). We next identified dorsal cells as cells expressing the *Iroquois Complex* genes *mirr*, *araucan* (*ara*) and/or *caupolican* (*caup*) (Fig S4). Triple-negative cells were identified as *v* cells (Fig 5A and Fig S3A’’,A’’’). Comparing control *v* cells from *mirr>ND-42^RNAi^* larvae with control cells from *mirr-Gal4* larvae by Differential Gene Expression (DGE) analysis revealed clear differences between these two populations of control cells (Fig S3C). Differences in Gene Ontology (GO) terms were associated with neuronal differentiation, possibly reflecting a sample-to-sample difference in staging (Fig S3D). We therefore used the *v* cells of the *mirr>ND-42^RNAi^* eye discs as internal controls to study the transcriptomic response to loss of Complex I activity. We identified 421 and 60 differentially expressed genes in *mirr>ND-42^RNAi^* UPs and DCs, respectively (Fig 5B,C; Fig S3E,E’; Table S3; most of these genes were up-regulated in *ND-42^RNAi^* cells). GO analysis showed that genes involved in ATP production and OxPhos were enriched in *ND-42^RNAi^* UPs (Fig 5E). Of note, most genes encoding glycolytic enzymes were found to be up-regulated in UPs (Fig 5D). Additionally, a glucose transporter (MFS3), a rate-limiting enzyme breaking glycogen into glucose (GlyP) and the rate-limiting enzyme contributing to the detoxification of a cytotoxic byproduct of glycolysis (GLO1) were also up-regulated (Table S3). In contrast, differentially expressed genes in DCs were predominantly associated with neuronal differentiation and morphogenesis (Fig S3E-F). Thus, increased glycolysis is a key metabolic response to cI inhibition. This response was primarily observed in UPs, possibly reflecting the metabolic constraints associated with the growth and proliferation of UPs.

**Figure 5:**
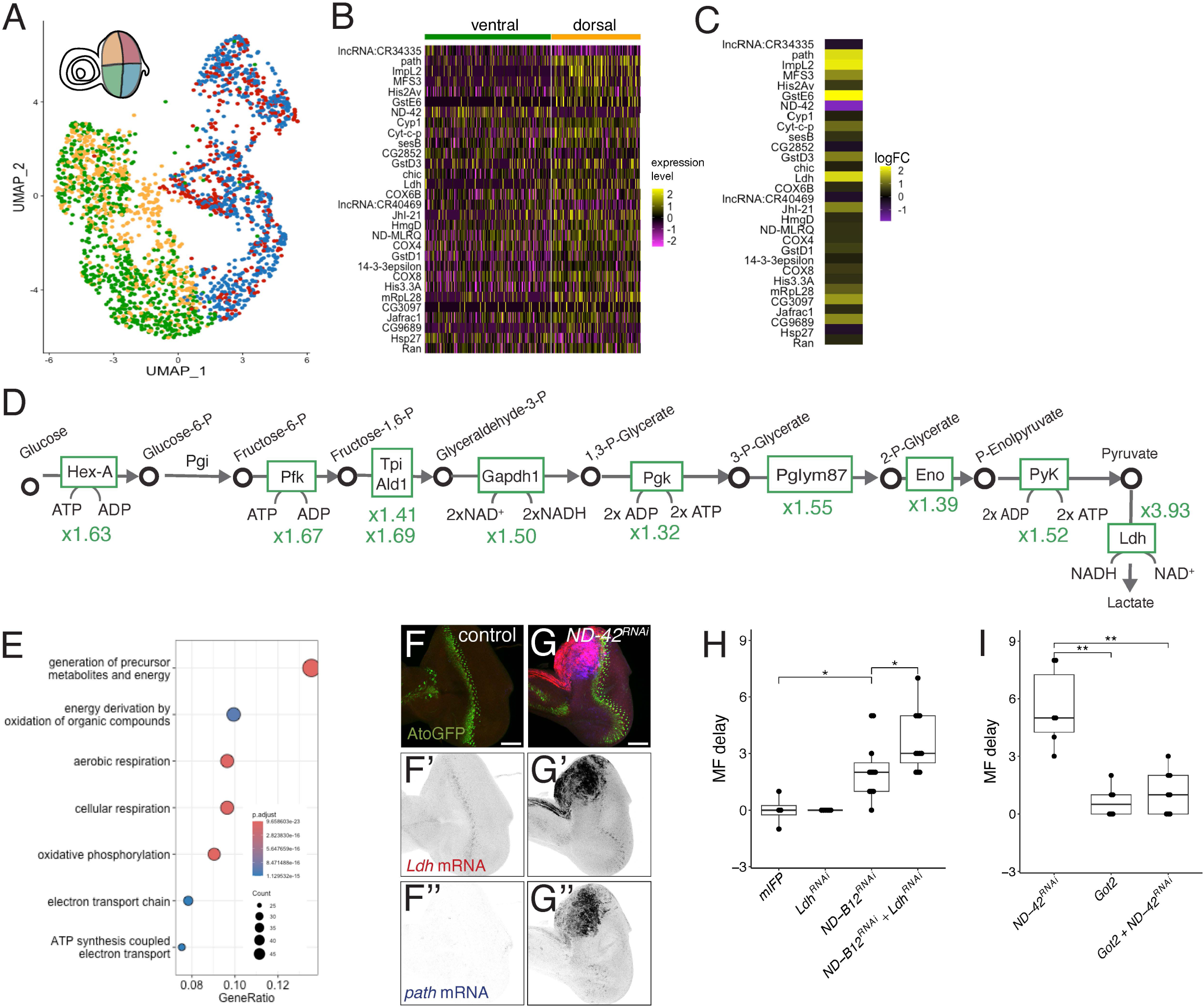
scRNAseq analysis of the compensatory response to ETC inhibition. **A)** UMAP analysis of *mirr>ND-42^RNAi^* eye field cells (color code as in the eye disc schematic). **B)** List of the top 30 differentially expressed genes between *d* (n= 528 cells) and control *v* UPs (n=747 cells) in *mirr>ND-42^RNAi^* eye discs. Genes were ranked by adjusted p-value. **C)** Relative changes in expression levels for the top 30 genes shown in B. The log2 Fold Change (logFC) values were color coded (see heat map). **D)** Glycolysis pathway with changes in gene expression (FC, green, between *d ND-42^RNAi^* and *v* control UPs). Increased glycolysis in *ND-42^RNAi^* cells was in part due to changes in gene expression. **E)** Gene Ontology (GO) term enrichment for biological pathways using the top 421 differentially expressed genes between *d* and *v* UPs in *mirr>ND-42^RNAi^* discs (adjusted p-value<0.05). **F-G’’)** The expression of the *Ldh* (red in F,G) and *path* genes (blue in F,G) was studied by smiFISH in control *mirrGal4 ato^GFP^* (F-F’’) and *mirr>ND-42^RNAi^ ato^GFP^* discs (G-G’’; AtoGPP, green, marked the differentiation front). The *Ldh* and *path* genes were up-regulated in UPs and dorsal peripodial cells upon silencing of the *ND-42* gene (G’,G’’). Note that the *Ldh* gene was expressed by a subset of MF cells in wild-type discs (F’), consistent with its proposed regulation *by* Ato (Aerts et al., 2010). **H)** The RNAi-mediated silencing of *Ldh* enhanced the MF delay resulting from the silencing of *ND-B12* (*Ldh^RNAi^,*0.0 +/−0.0, n=6; *ND-B12^RNAi^*, 2.2 +/−1.6, n=11; *ND-B12^RNAi^* + *Ldh^RNAi^*, 4.1 +/− 2.4, n=12). The mIFP data shown in Fig 1E were used as negative control in H. **I)** Overexpression of Got2 enhanced the MF delay resulting from the silencing of *ND-42* (*Got2*, 0.7 +/−0.8, n=6; *ND-*42*^RNAi^*, 5.5 +/−2.1, n=6; *ND-42^RNAi^* +*Got2*, 0.7 +/−0.9, n=6). Wilcoxon tests: * < 0.05; ** < 0.01. Scale bar, 50µm.

Increased glycolysis is often accompanied by an increased rate of lactate production from pyruvate catalyzed by Ldh because this reaction regenerates NAD^+^ from NADH and NAD^+^ is required to fuel glycolysis and (Luengo et al., 2021). Consistent with increased glycolysis in UPs, we observed increased *Ldh* gene expression in UPs by scRNAseq (Fig 5D) and smiFISH (Fig 5F-G’). We therefore hypothesized that increased Ldh activity promotes glycolysis via the regeneration of NAD^+^. To test whether increased Ldh was part of a compensatory response to cI inhibition, we silenced the *Ldh* gene in combination with *ND-B12*, another cI subunit gene (Fig 5H, Table S1; double silencing of the *Ldh* and *ND-42* genes was not technically feasible because both RNAi constructs are at the same genomic site). While the RNAi-mediated silencing of *Ldh* had no effect on furrow progression, it strongly enhanced the weak *ND-B12^RNAi^* MF delay phenotype (Fig 5H). Thus, increased Ldh activity appeared to compensate for the loss of cI activity. We propose that Ldh acts in this context by regenerating NAD^+^ from NADH, allowing for increased glycolysis and compensatory ATP production. Metabolomic analysis of eye-antenna discs by LC/MS further supported the notion that NAD^+^ could be limiting upon cI inhibition (Fig S5). Indeed, the accumulation of Glyceraldehyde-3-phosphate (GAP) indicated that the NAD^+^-dependent reaction using GAP as a substrate becomes rate-limiting; the accumulation of Glycerol-3-phosphaste (G3P) and Dihydroxyacetone phosphate (DHAP) that are produced from fructose-1,6-diphosphate (FDP) and GAP suggested that NAD^+^-producing reactions are enhanced and that the Glycerol Phosphate shuttle is activated to regenerate NAD^+^ in the cytoplasm (Fig S5C,D). Together, these data indicated that the rate of NAD^+^ regeneration could be limiting glycolysis upon cI inhibition.

Beyond glycolysis, amino acid import also appeared to be up-regulated upon ETC inhibition. Two transporters known to support growth upon nutrient restriction (Feng et al., 2020; Rebelo and Homem, 2023), Pathetic (*path*) and Jhl-21 were also up-regulated in UPs upon silencing of *ND-42* (Fig 5B,C,F’’-G’’ and Table S3). These gene expression changes appeared to also reflect a compensatory response since the RNAi-mediated silencing of *path* enhanced the *ND-42* MF delay phenotype (data not shown). Previous studies have shown that Aspartate is limiting for cell growth upon cI inhibition in mammals, and that the increased NADH/NAD+ ratio decreases the ability of cells to synthesize Aspartate, a precursor for the synthesis of Glutamate and Proline (Birsoy et al., 2015; Sullivan et al., 2015). To test whether Aspartate was also limiting in eye cells upon cI inhibition, the Glutamate oxaloacetate transaminase 2 (Got2) was overexpressed to favor Aspartate synthesis from OxaloAcetate. Increased Got2 activity was found to suppress the *ND-42^RNAi^* MF delay phenotype (Fig 5I). Thus, Aspartate appeared to be limiting upon cI inhibition for the timely progression of the MF in *Drosophila*. We therefore propose that increased amino acid import compensated for reduced amino acid biosynthesis upon ETC inhibition.

Our analysis also identified mitochondria biogenesis and/or activity genes, and several detoxification genes (*GstD1*, *GstD3*, *GstE1*, *GstE6, GstE8, GstT1, Nmdmc*) (Sharma et al., 2004; Yu et al., 2015) were up-regulated upon the silencing of *ND-42* (Table S3) These data were consistent with increased levels of Radical Oxygen Species (ROS) upon loss of Complex I activity (Owusu-Ansah et al., 2013). We also noted that a secreted antagonist of Insulin signaling (ImpL2), which mediates tissue wasting (Figueroa-Clarevega and Bilder, 2015), and the relaxin-like hormone dIlp8 were expressed at higher levels upon loss of cI activity. This suggested that a local perturbation in cellular metabolism in the eye triggered a systemic response to slow down tissue growth and developmental progression at the organismal level. This systemic response likely contributed to the developmental delay noted in *mirr>ND-42^RNAi^* larvae (see also Fig 3D).

Finally, we noted that several transposable elements (TEs) were expressed at a higher level in UPs and DCs upon cI inhibition (Fig S3G,H and Table S4). Since increased glycolysis has broad impact on histone modifications (Dai et al., 2020) and since TEs are usually silenced epigenetically, we speculate that these changes in TE expression might reflect a global impact of the cellular metabolism on the epigenome. Consistent with this possibility, cI inhibiton led to increased level of Acetyl-CoA and S-Adenosyl Methionine (SAM) that act as donor groups for histone acetylation and methylation (Fig S5D).

In summary, our analysis showed that *ND-42^RNAi^* cells elicited both local and systemic responses to loss of ETC activity, and that increased glycolysis was a key compensatory response elicited by the proliferative undifferentiated cells, which likely resulted in a strong demand for NAD^+^(Luengo et al., 2021).

### The speed of the MF depends on the metabolic state of the undifferentiated cells

Our data above showed that metabolic rewiring was particularly noticeable in UPs. This compensatory response to cI inhibition was consistent with the observation that *ND-42^RNAi^* UPs were still actively growing and proliferating despite cI inhibition. This observation suggested that the speed of progression of the MF was dependent on the metabolic state of these cells. To test this, we sought to restrict the RNAi-mediated silencing of *ND-42* to UPs using a RNAi-resistant form of ND-42 expressed specifically in cells posterior to the MF. We therefore designed a RNAi-resistant *ND-42* gene, noted *ND-42\**, by recoding the region of its Open Reading Frame targeted by the RNAi construct (see Methods). The expression of *ND-42\** under the control of a constitutive promoter was sufficient to restore the progression of the MF in *mirr>ND-42^RNAi^* larvae (Fig 6A), hence validating this approach. We then used strong enhancers from the *atonal* (*ato*) gene to express *ND-42\** in MF cells and examined whether these *ato-ND-42\** transgenes could restore OxPhos activity in the *mirr>ND-42^RNAi^* cells posterior to the MF. Using the iATPsn sensor to examine changes in ATP levels upon inhibition of the ETC by AntA, we found that expression of ND-42* in the MF restored production of ATP by the ETC in *mirr>ND-42^RNAi^*DCs (Fig 6B). We conclude that the ND-42* protein restored OxPhos activity upon silencing of the endogenous *ND-42* gene and that the ND-42* protein was stable enough to rescue cI activity in all DCs. Analysis of the iATPsn fluorescence signal along the AP axis showed that expression of ND-42* in the MF rapidly restored OxPhos activity in DCs (Fig 6C,D). Thus, RNAi-mediated silencing appeared to be restricted to cells located anterior to, and at the MFs in these discs. We next examined the MF delay in these discs and found that restoring ND-42 activity in cells posterior to the MF did not suppress the delay in MF progression (Fig 6A). This indicated that the activity of ND-42 was required in UPs and/or MF cells for the timely progression of the MF. Finally, to test whether the metabolic state of the DCs may also be required to set the the speed of the MF, we sought to specifically perturb the metabolic state of the DCs. To do so, we restricted the expression of a GFP-tagged version of Foxo, which was found to delay the progression of the MF when expressed in all *d* cells (Fig 6E,G), in DCs using a strong enhancer from the *E(spl)m4-BFM* gene which is active in the MF (Couturier et al., 2024), and then used *mirr>GFP^RNAi^* to silence FoxoGFP expression in *d* cells, hence restrict the expression of FoxoGFP to *v* cells. In this context, no MF delay was observed in *v* cells (Fig 6F,G), suggesting that the metabolic state of the DCs does not play a critical role in the progression speed of the MF. We therefore conclude that the metabolic states of the UPs and/or MF cells determine the speed of progression of the MF.

**Figure 6:**
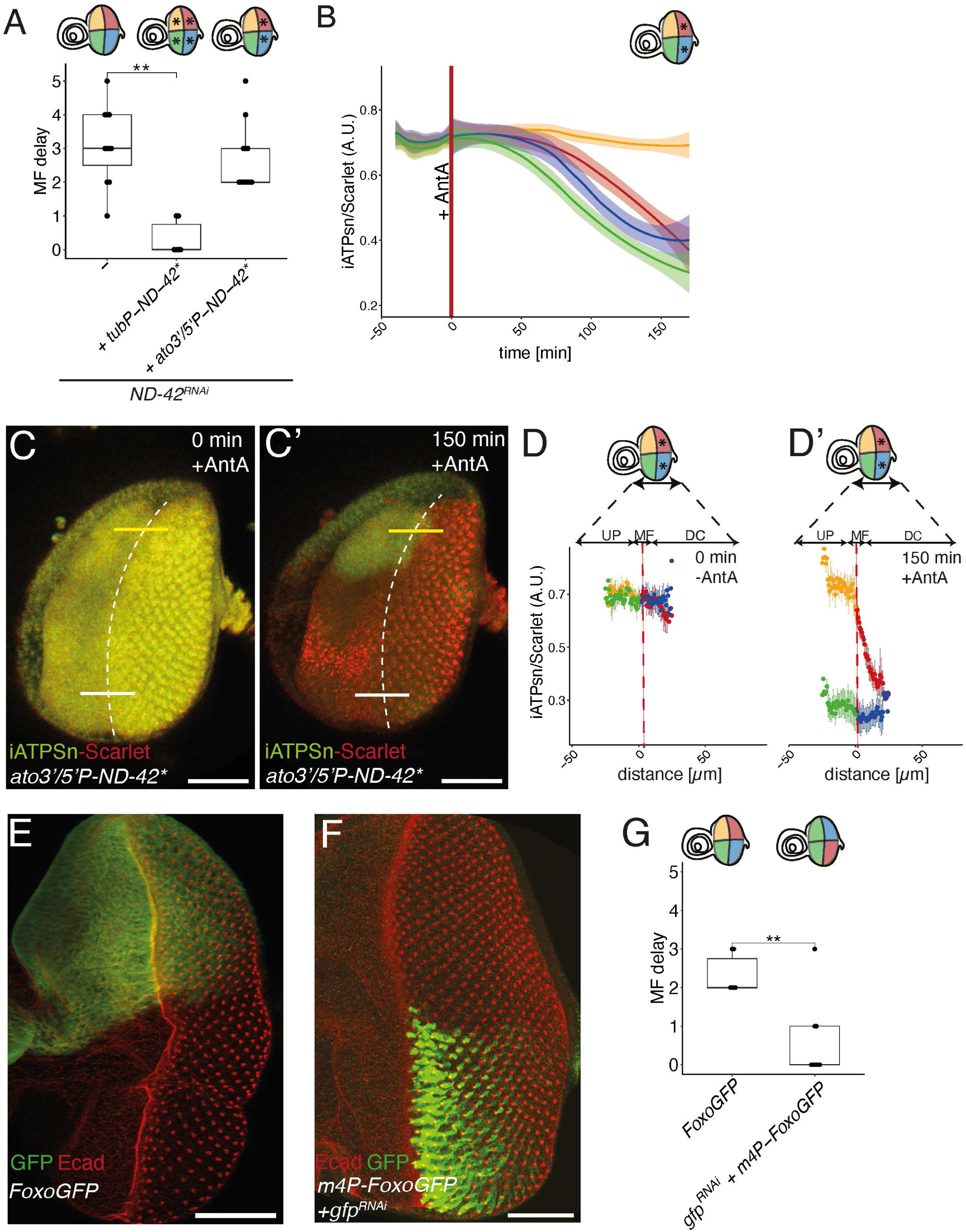
A UP-specific metabolic perturbation slowed down the progression of the front. **A)** A UP-specific perturbation in energy metabolism was produced in *mirr>ND-42^RNAi^* discs by expressing a RNAi-resistant form of ND-42, ND-42*, in DCs only (as indicated with * in DC quadrants). As a positive control, ND-42* rescued the *ND-42^RNAi^* MF delay phenotype when expressed ubiquitously (as indicated with * in all four quadrants of the eye). This UP-specific perturbation resulted in a strong MF delay: *mirr>ND-42^RNAi^*, 3.9 +/−1.1 (n=11); *mirr>ND-42^RNAi^ tubP-ND-42\**, 0.3 +/−0.5 (n=6); *mirr>ND-42^RNAi^ ato3’/5’P-ND-42\**, 2.6 +/−0.9 (n=12). **B-D’)** UP-specific perturbation in *mirr>ND-42^RNAi^ ato3’/5’P-ND-42\** eye discs. Cells with increased glycolysis were identified using the iATPsn sensor based on their response to AntA. Time course analysis of the normalized iATPSn signal showed that the lack of response to AntA, indicative of increased glycolysis, was restricted to dorsal UPs (B; n=4; the data are shown as in Fig 4M). The iATPSn signal (green; Scarlet, red) is shown on snapshot views of a disc at t=0 (C) and t=150 (C’). This indicated that the loss of endogenous *ND-42* activity was efficiently compensated by ectopic ND-42*. The normalized iATPSn signal was plotted along the AP axis (yellow line on *d* side and white line for *v* side in C,C’) at t=0 (D) and t=150 (D’; n=4). This showed that the suppression of the *ND-42^RNAi^*phenotype was first detected at the MF. **E-G)** Expression of FoxoGFP in *mirr>FoxoGFP* discs (E) led to a MF delay phenotype (G; 2.3 +/−0.5, n=6). In contrast, restricting the expression of FoxoGFP to ventral DCs in *m4P-FoxoGFP mirr>gfp^RNAi^* discs (F) had no effect on MF progression (0.6 +/−1.0, n=9). This indicated that restricting metabolic perturbations to DCs did not slow the progression of the MF. Wilcoxon tests: ** < 0.01. Scale bar, 50µm.

### Cellular NAD^+^ availability appeared to constrain developmental speed

We next examined whether the rate of NAD^+^ regeneration from NADH could be limiting the speed of progression of the MF upon cI inhibition. To test this, we over-expressed a bacterial NADH oxidase, LbNOX, to increase the rate of NAD^+^ regeneration (Titov et al., 2016). Expression of LbNOX in the cytosol, using cyto-LbNOX, had no detectable effect on MF progression (Fig 7A) and only slightly decreased the NADH/NAD^+^ ratio in differentiated eye disc cells (Fig 7I; the weak effect measured in UPs was not statistically significative).

**Figure 7:**
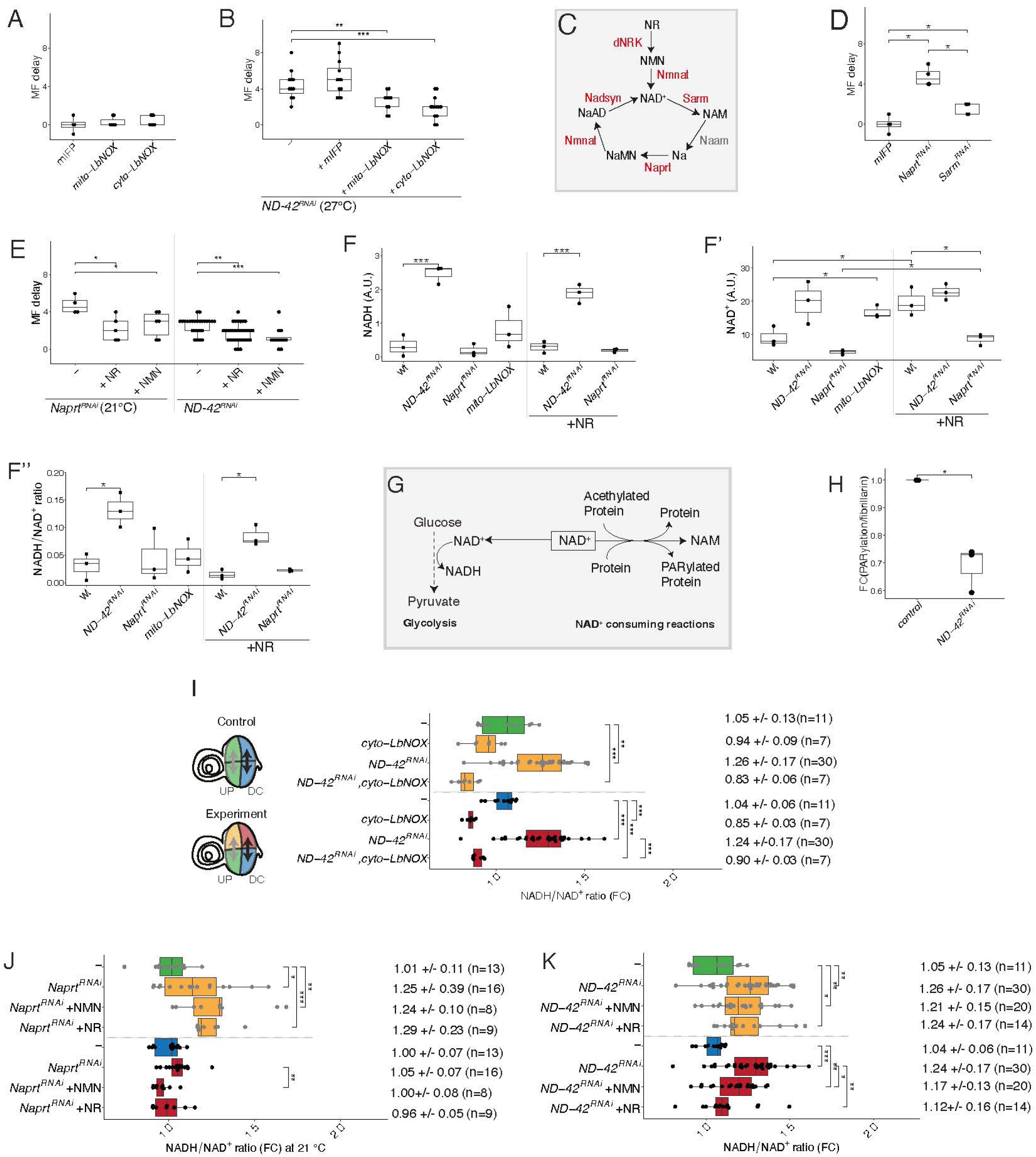
The rate of NAD^+^ regeneration appeared to limit developmental speed. **A,B)** MF delay analysis showing that expression of cyto-LbNOX and mito-LbNOX had no significant effect on MF progression (A; control mIFP data are from Fig. 1E) but efficiently suppressed the *ND-42^RNAi^* MF delay phenotype (B). The strong MF delay seen in *mirr>ND-42^RNAi^* at 27°C (4.4 +/−1.6, n=11; negative control with mIFP co-expression: 5.3 +/−2.0, n=12) was suppressed by mito-LbNOX (2.6 +/−1.1, n=10) and cyto-LbNOX (1.9 +/−1.3, n=13). **C)** A simplified view of NAD^+^ metabolic pathway: NAM, nicotinamide; Na, Nicotinic acid; NaMN, Nicotinic acid MonoNucleotide; NaAD, Nicotinic acid Adenine Dinucleotide. The genes tested for possible MF delay by RNAi-mediated silencing and/or over-expression are indicated in red (C). **D)** Silencing the *Sarm* and *Naprt* genes delayed the progression of the MF (*Sarm^RNAi^:* 1.6 +/− 0.5, n=5; *Naprt^RNAi^*: 4.8 +/−1.0, n=4). **E)** The *Naprt^RNAi^* phenotype was suppressed in part by the addition of NR (2.2 +/−1.3, n=5) and NMN (2.6 +/−1.4, n=6). Likewise, the *ND-42^RNAi^* defect (2.6 +/−0.9, n=25) was partially suppressed by adding NR (1.8 +/−1.0, n=33) or NMN (1.25 +/−1.1, n=10). **F)** LC/MS analysis of NAD^+^ and NADH in whole cell lysates from control *tub^ts^> +*, *tub^ts^*>*ND-42^RNAi^*, *tub^ts^*>*Naprt^RNAi^*and *tub^ts^*>*mito-LbNOX* eye-antenna discs (n=3 for all genotypes). Addition of NR restored NAD^+^ levels in *Naprt^RNAi^* eye-antenna discs (the same control *tub^ts^*>*ND-42^RNAi^* data appear in Fig 2 K-K’’). Note that mito-LbNOX expression appeared to increase NAD^+^ levels relative to control eye disc. Statistical analysis was performed between control and all experimental genotypes, separately for the two food conditions (+/− NR), and between food conditions for each genotype. **G)** NAD^+^ has two functions, as a redox cofactor in energy metabolism, e.g. glycolysis, and as a co-substrate in NAD^+^-consuming reactions. For instance, Parp1 and Sirtuins use NAD^+^ to poly-PARylate (blue) or deacetylate (red) proteins. **H)** The normalized levels of poly-PARylation levels were measured in *ND-42^RNAi^*(*tub^ts^>ND-42^RNAi^*), and *pdhb^RNAi^*(*tub^ts^>pdhb^RNAi^*) relative to control (*tub^ts^>+*), eye discs using Western Blot analysis (the Fibrillarin signal was used for normalization). Reduced poly-PARylation was observed in *ND-42^RNAi^* and *pdhb^RNAi^*discs. **I)** Analysis of the effect of cyto-LbNox on the NADH/NAD^+^ ratio (FC of the Peredox values measured in *d* and *v* cells plotted as in Fig 4I). In all experiments, *d* cells were identified using the infra-red signal produced by mIFP (transgene present in the four genotypes studied here). Data from control UPs shown and DCs are in green and blue, respectively, whereas experimentally perturbed UPs and DCs are in orange and red, respectively (see eye schematics). The expression of cyto-LbNOX in *mirr>cyto-LbNox*+*mIFP* discs suppressed the effect of *ND-42^RNAi^* (control and *ND-42^RNAi^*data were from Fig 4I). **J)** Analysis of the effect of *Naprt^RNAi^* on the NADH/NAD^+^ ratio measured as in G (this experiment was performed at 21°C to improve the viability of *Naprt^RNAi^* larvae). Inhibition of the NAD^+^ recycling pathway led to elevated NADH/NAD^+^ ratio in UPs but not DCs. Adding NR and NMN did not change the NADH/NAD^+^ ratio measured in UPs. **K)** Analysis of the effect of NAD^+^ precursors on the NADH/NAD^+^ ratio measured in *mirr>ND-42^RNAi^* discs using Peredox (as in panel F). The addition of NR and NMN had no effect on the NADH/NAD^+^ ratio in UPs and reduced this ratio only in DCs. Note that the control (*mirr>mIFP*) and *ND-42^RNAi^* data were from Fig 4I. Wilcoxon tests in A-E and H-K: *< 0.05; **< 0.01; ***< 0.0001. 2-sided t-tests in F-F’’: * < 0.05; ** < 0.01.

Likewise, expression of LbNOX in mitochondria, using mito-LbNOX, did not delay the MF and had no significant effect on the NADH/NAD^+^ ratio as measured by LCMS in eye-antenna disc extracts (Fig 7F’’). However, the levels of both NAD^+^ and NADH appeared to be increased upon mito-LbNOX expression (Fig 7F,F’), suggesting that increasing the pools of NAD^+^ might compensate for a loss of reduced NADH in mitochondria.

We next combined cyto-LbNOX and mito-LbNOX with *ND-42^RNAi^* and found that both cyto-LbNOX and mito-LbNOX were sufficient to partially suppress the MF delay seen in *mirr>ND-42^RNAi^* discs (Fig 7B). A similar effect was observed in *mirr>ND-49^RNAi^* discs (Fig S6A; *ND-49* encodes another cI subunit gene; see Table S1). This showed that increasing the rate of NAD^+^ regeneration partially restored developmental speed in cI-inhibited eye discs. Peredox analysis further showed that expression of cyto-LbNOX fully suppressed the effect of *ND-42^RNAi^* on the NADH/NAD^+^ ratio (Fig 7I). Thus, the rate of NAD^+^ regeneration from NADH appeared to be limiting upon cI inhibition for the timely progression of the MF.

To further examine the role of NAD^+^ in the progression speed of the MF, we altered the level of cellular NAD^+^ by adding precursors of NAD^+^ biosynthesis in the fly food and/or by silencing genes of the NAD^+^ salvage pathway (Gossmann et al., 2012) ( Fig 7C). Testing five of the six enzymes involved in the NAD^+^ salvage pathway, we found that silencing the *Sterile alpha and Armadillo motif* (*Sarm*) and *Nicotinate phosphoribosyltransferase* (*Naprt*) genes led to a MF delay phenotype (Fig 7D, Fig S6B). Analysis of *Ldh* gene expression in *mirr>Naprt^RNAi^* eye discs showed that Ldh expression was increased in UPs (Fig S6C,C’), indicating that NAD^+^ was limiting upon loss of *Naprt* activity. Consistently, adding the NAD^+^ precursors Nicotinamide Riboside (NR) and Nicotinamide MonoNucleotide (NMN) to the fly food increased the level of NAD^+^ in *Naprt^RNAi^* eye-antenna disc cells (Fig 7F’) and suppressed the *Naprt^RNAi^* MF delay phenotype (Fig 7E). We therefore conclude that limiting NAD^+^ availability in *mirr>Naprt^RNAi^* discs resulted in a MF delay. To further test whether NAD^+^ could also limit developmental speed in *ND-42^RNAi^* cells, we added NR or NMN to the food of mirr>*ND-42^RNAi^* larvae. Adding these NAD^+^ precursors led to a partial suppression of the *ND-42^RNAi^* MF delay phenotype (Fig 7E). This suggested that NAD^+^ constrained the speed of progression of the MF upon cI inhibition.

To further test whether free NAD^+^ was limiting upon cI inhibition, we examined the activity of a non-redox NAD^+^-dependent enzyme. Indeed, beyond its role as a co-factor in many redox reactions, including glycolysis, NAD^+^ is also an essential co-substrate for non-redox NAD^+^-dependent enzymes, including Sirtuins and poly(ADP-ribose) polymerases (PARPs; Fig 7G, Fig S6E). Since protein poly-PARylation by Parp1 consumes NAD^+^, lowering the level of free NAD^+^ is expected to decrease protein PARylation. To test whether free NAD^+^ is limiting in *ND-42^RNAi^* cells, we measured the levels of poly-PARylated proteins in whole disc extract of *tub^ts^> ND-42^RNAi^* eye-antenna discs by Western Blots analysis. A strong decrease in protein poly-PARylation was observed upon cI inhibtion (Fig 7H). This further indicated that NAD⁺ is limiting upon cI inhibition, likely reflecting a strong demand for NAD⁺ by glycolysis.

Studies in mouse cultured cells showed that supplementation of NAD^+^ precursors not only increased the pool of NAD+ but also changed the NADH/NAD+ ratio (Figley et al., 2021; Hu et al., 2021). Likewise, we found that adding NR not only increased the level of NAD^+^ in control eye-antenna discs (Fig 7 F,F’) but also reduced the NADH/NAD^+^ ratio in control UPs (Fig S6D; a similar effect was seen with NMN addition; however, no significant change in NADH/NAD^+^ was detected in whole discs by LC/MS, Fig 7F’’). Conversely, down-regulating NAD^+^ levels through the inhibition of the salvage pathway in *mirr>Naprt^RNAi^* discs was also associated with an increase in the NADH/NAD^+^ ratio in UPs (Fig 7J). Additionally, we observed above that changing the redox state by expressing cyto-LbNOX led to increased NAD^+^ levels (Fig 7F,F’). Thus, NADH/NAD^+^ ratio values and cellular NAD^+^ levels appeared to be inter-dependent. Despite this, we observed that the suppression of the *Naprt^RNAi^* MF delay by the addition of NR and NMN was not accompanied by detectable changes in the NADH/NAD^+^ ratio in UPs (Fig 7J). Likewise, the suppression of the *ND-42^RNAi^* MF delay phenotype by NR and NMN was not associated with changes in the NADH/NAD^+^ ratio in UPs (Fig 7K). Since a change in the progression of the MF was not associated with a detectable change in cellular redox in UPs, this suggested that limiting NAD^+^ appeared to constrain the speed of progression of the MF in the developing eye disc of *Drosophila*.

## Discussion

The eye imaginal discs of *Drosophila* offers two key advantages to study *in vivo* the speed of developmental patterning. First, the crystal-like pattern of ommatidia that forms behind the differentiation front permits to easily map time onto space. Second, gene-specific perturbations can be restricted to the dorsal compartment of the developing eye disc, allowing us to study relative changes in developmental speed at the tissue level independently of any associated changes taking place at the organismal level. Here, we made use of these two advantages to identify via a simple assay specific perturbations that are associated with a relative change in the progression speed of the differentiation front. The importance of these advantages can be illustrated with the *ND-42^RNAi^* perturbation that induced a developmental delay associated with the up-regulation of dilp8, which is known to reduce the production of the steroid hormone ecdysone, yet produced an easy readout in the MF delay assay. A well-known limitation of RNAi is that this approach often results in partial loss of function (Dietzl et al., 2007). This limitation actually turned out to be useful for uncovering the role of key metabolic proteins that have essential functions. Moreover, silencing efficiency could be further modulated using the Gal4/Gal80ts system. For instance, while a delay in MF progression was detectable upon conditional silencing of the translation initiation factor eIF2α, a stronger down-regulation led to defective eye growth. Finally, the issue of specificity, or off-targets, could be addressed using more than one RNAi line per gene, or by targeting different subunits of protein complexes.

Loss of function perturbations directly affecting the mitochondrial metabolism were found to slow down the progression of the furrow. Thus, our work extends to insects the recent conclusion that ETC activity and mitochondrial activity regulates the rate of development in mammals (Diaz-Cuadros et al., 2023; Iwata et al., 2023). The inhibition of the ETC activity could drive metabolism away from its optimum, leading to inefficient energy consumption, with more energy being spent at maintaining structures, possibly associated with increased rate of ATP production (Sturm et al., 2023) and less energy devoted to energetically costly processes, e.g. the synthesis of proteins with rapid turn-over. In the *Drosophila* eye, ETC dysfunction should lead to a loss of mitochondrial ATP production which appeared to be compensated by an increase in glycolysis. Gene expression studies indicated that glycolysis and amino acid import were predominantly up regulated in UPs anterior to the MF, possibly reflecting a higher demand for energy associated with growth and proliferation. Indeed, the costs associated with growth and division, including the cost of protein synthesis (Buttgereit and Brand, 1995), is thought to be one order of magnitude higher than the costs associated with maintenance (Lynch and Marinov, 2015). The loss of ETC activity in eye cells also led to the accumulation of NADH. Increased NADH levels relative to NAD^+^ were seen not only in mitochondria but also in the cytosol and nucleus. High NADH were proposed to have detrimental effects on the cellular metabolism and previous studies had shown that these can be alleviated by enhancing the regeneration of NAD^+^ via enhanced production of lactate by Ldh (King and Attardi, 1989) or via the expression of the bacterial NADH oxidase LbNOX (Titov et al., 2016). Consistent with this, expression of LbNox in mitochondria or in the cytosol suppressed in part the effect of *ND-42^RNAi^*and Ldh was over-expressed in the *ND-42^RNAi^* eye UP cells that grow and proliferate. Moreover, while Ldh is expressed at low levels and its activity is largely dispensable in control eye cells, the activity of Ldh became critical upon cI inhibition. Thus, excessive NADH, limiting NAD^+^ and/or elevated NADH/NAD^+^ appeared to have detrimental effects on cellular metabolism and were associated with slow developmental speed in the *Drosophila* eye.

A recent study showed that the NADH/NAD^+^ ratio plays a causal role in the regulation of developmental speed in mammals, with slower developmental speed in human cells than in mouse cells caused by a higher NADH/NAD^+^ ratio (Diaz-Cuadros et al., 2023). Most of our data are consistent with this view. Notably, an elevated cellular NADH/NAD^+^ ratio correlated with a slow progression of the MF upon *ND-42* or *Naprt* knock-down. However, we also noted that an increased NADH/NAD^+^ ratio positively correlated with a high demand for NAD^+^. For instance, limiting NAD^+^ availability was associated with an increased NADH/NAD^+^ratio upon inhibition of the NAD^+^ recycling pathway. Conversely, increasing NAD^+^ availability by NR addition reduced the NADH/NAD^+^ ratio in growing UP cells of control larvae. Likewise, decreasing the NADH/NAD^+^ ratio by expressing mito-LbNOX led to a compensatory increase in the level of NAD^+^ (this compensatory response could help maintain a normal redox state). Also, increased NADH/NAD^+^ in *ND-42^RNAi^* cells appeared to be associated with limiting NAD^+^, as suggested by i) the suppression of the *ND-42^RNAi^* MF delay phenotype by NR; ii) the decreased protein PARylation measured in *ND-42^RNAi^* cells; iii) our transcriptomic and metabolomic results indicating that reactions regenerating NAD^+^ from NADH were up-regulated (see for instance the increased expression of the *Ldh* gene), whereas reactions producing NADH from NAD^+^ were inhibited (as suggested for instance by the increased accumulation of GAP in the glycolysis pathway). These observations strongly suggested that a strong inter-dependency exists between changes in cellular redox and NAD^+^ availability. This in turn implied that disentangling the effects due to changes in NADH/NAD^+^ ratio from those due to reduced NAD^+^ availability may prove to be a complex task. Despite this, two observations suggested that limiting NAD^+^ availability had an impact beyond its effect on cellular redox on the speed of progression of the MF in the fly eye. First, addition of NAD^+^ precursors restored in part the speed of progression of the MF in *mirr>Naprt^RNAi^* cells without changing the NADH/NAD^+^ ratio measured in UP cells. Second, addition of NAD^+^ precursors partly suppressed the *mirr>ND-42^RNAi^* MF delay phenotype, indicative of restored developmental speed, whereas the cellular redox of UPs remained largely unchanged. Since the metabolic state of UPs determined the speed of MF progression, this suggested that a change in developmental speed may not necessarily correlate with a change in cellular redox. We therefore suggest that the availability of NAD^+^ contributes to limit the speed of development in the *Drosophila* eye. Interestingly, NAD^+^ is not only a key redox cofactor for energy metabolism but is also a substrate for protein post-translational modifications enzymes. Modifications include ribosylation and acetylation, and protein targets includes glycolytic enzymes, histones and microtubules (Kulkarni and Brookes, 2019). Therefore, NAD^+^ couples energy metabolism with gene expression and cellular organization, and maintaining proper NAD^+^ levels is key in various physiological contexts in mammals (Abdellatif et al., 2022; Beltrà et al., 2023; Brenner, 2022; Hosios and Vander Heiden, 2018; Luengo et al., 2021; Romani et al., 2021; Zhang et al., 2024). In *Drosophila* NAD^+^ levels vary with temperature, with lower levels observed at low and high temperature, and higher levels at optimal temperature for adult physiology and lifespan, therefore suggesting that low NAD^+^ may reflect a departure from optimal cellular homeostasis (Klepsatel et al., 2020). Future work will address whether reduced NAD^+^ availability may limit the activities of enzymes that depend on NAD^+^ as a cofactor or use NAD+ as a co-substrate, and whether the resulting metabolic perturbations affect developmental speed in the *Drosophila* eye.

## Acknowledgements

We thank S. Aerts (VIB, Leuven), M. Baylies (MSKCC, New-York), B Hassan (ICM, Paris), J.H. Camacho (IP, Paris), L. Cantini (IP, Paris), J. Knoblich (IMBA, Vienna), P. Leopold (IC, Paris), J. Marvin (JRC, HHMI), M. Miura (Univ. of Tokyo), L. J. Neukomm (Univ. of Lausanne), J. Rutter (Univ. of Utah), T. Ryan (Cornell Univ.), Cyril Scandola (SEM facility, IP, Paris), G. A. Smith (Cardiff Univ.), N. Tapon (Crick Inst., London), E. Vimont (IP, Paris), Y. Zhao (IP, Paris), Flybase, FlyORF, the Vienna Drosophila Ressource Center (VDRC) and the Bloomington Drosophila Stock Center (BDSC) for flies, plasmids, antibodies, database services, technical assistance and/or scientific guidance. We thank M. Rera, B. Hassan and D. Green for critical reading. NV and YG were supported by the Pasteur Paris University (PPU) International Doctoral Program. NV and CP received support from the Labex Revive (ANR-10-LABX-0073). YG was supported by the Chinese Biotechnology Company (CNBG). This work was funded by grants from the Agence Nationale pour la Recherche to FS (ANR-10-LABX-0073 and ANR-22-CE13-0016-01).

## Methods

### Flies

A *mirr-Gal4* driver, *mirr^DE^*, was used to drive gene-based perturbations in *d* cells. To easily mark the differentiation front, *ato^GFP^* was recombined with *mirr^DE^*(*ato^GFP^*is a functional GFP knock-in allele of *ato*). Unless noted otherwise, all crosses and flies were maintained at 25°C for analysis. Conditional silencing was achieved at 29°C for 3 days using a *mirr-Gal4 tubP-Gal80^ts^ ato^GFP^* driver for MF delay analysis, and a *tubP-Gal4 tubP-Gal80^ts^* driver for NAD^+^/NADH-glo assay analysis as well as for the LC/MS experiments. NMN (β-Nicotinamide mononucleotide, N3501, Sigma) and NR (Nicotinamide riboside chloride, SMB00907, Sigma) were dissolved in water and added to the fly food (NMN, 25 mM; NR, 5 mM) before introducing first (L1) and second (L2) instar larvae. After 3-4 days at 25 °C, mid-L3 larvae were collected for analysis. The *UAS-mIFP-T2A-HO1* transgene (BL-64181) that encodes an infrared marker was recombined with *mirr^DE^*, Peredox sensor and SoNar sensor analysis was performed using this driver line.

The speed of progression of the MF was measured in eye discs dissected from *hs-flp*/+;; *mirr-Gal4 ato5’-FRT-mCherrynls-FRT-GFPnls*/+ third instar larvae (L3) which were subjected to a mild heat-shock (hs; 36°C for 30 min) to induce flp-out recombination and then dissected after a chase period of 4 to 16 hr post-hs.

The following transgenic lines were obtained by phiC31-mediated integration: PBac{y[+], pUbi-Peredox-CherryNLS attB-3B}VK31 (62E1); PBac{y[+], pUbi-ATPSnFR-95A.119L-mpHmScarlet attP}VK31 (62E1); PBac{y[+], pUAS-cyto-LbNOX attP-3B}VK37 (22A3); PBac{y[+], pUAS-mito-LbNOX attP-3B}VK37 (22A3); PBac{y[+], ato5’-FRT-mCherrynls-FRT-GFPnls attB-3B}VK31 (62E1). Injection was performed by BestGene Inc. (Chinmo, USA). Flies were grown on a rich medium: corn flour (60g/L), yeast extract (60g/L), agar (6g/L), nipagin (4g/L), propionic acid (0.4%).

### Transgenes

The nuclear Peredox NADH/NAD^+^ ratio sensor was obtained by cloning a PCR-amplified fragment encoding the Peredox-mCherry-NLS sensor (Addgene #3238) into a pUbi-attB vector using Gibson Assembly. The Cyto-Sonar sensor was obtained by cloning a PCR amplified fragment encoding SoNar obtained from the genome of UAS-SoNar flies (Bonnay et al., 2020) into a pUbi-attB vector using Gibson Assembly. The mito-SoNar sensor was similarly obtained, adding 2 N-terminal copies of the mitochondria targeting signal from the human *cox8A* gene and a GDPPVAT linker which were obtained by the annealing of two partially overlapping oligonucleotides followed by PCR amplification. The ATP sensor was similarly obtained using a PCR-amplified fragment encoding the iATPsnFR-A95A.A119L.mpHmScarlet sensor (Marvin et al., 2024). A 1986 nucleotide (nt)-long fragment starting 5’ to the ATG of the *Ubi-p63E* gene (*CG11624*) and encoding its promoter was used to direct the constitutive expression of a Peredox-mCherrynls fusion protein.

The UAS-cyto-LbNOX plasmid was obtained by cloning a PCR-amplified fragment encoding cyto-LbNOX (Addgene #7528) into a pUAS-attB vector using Gibson Assembly. A short DNA fragment encoding one copy mitochondria targeting signal from the human *cox8A* gene and a GDPPVAT linker was obtained via the annealing of two partially overlapping oligonucleotides followed by PCR amplification and was cloned into the UAS-cyto-LbNOX plasmid by Gibson assembly to generate the UAS-mito-LbNOX transgene.

To measure the speed of progression of the front, we designed a recombination-based Cherry-to-GFP switch plasmid, noted ato5-FRT-mCherrynls-FRT-GFPnls. This plasmid, produced in three Gibson assembly steps, consists in the ato5’Eye enhancer of the *atonal* (*ato*) gene (a 2918 nt fragment located 7471 nt 5’ to the *ato* start codon), the *ato* basal promoter (a 1058 nt fragment located 58 nt 5’ to the *ato* start codon), a FRT-mCherrynls-hsp70 stop-FRT cassette followed by a GFPnls obtained from pattB-neur-GFPnls-SV403’UTR plasmid (Aerts et al., 2010).

A RNAi-resistant ND-42 gene, ND-42*, was obtained by recoding the 300nt fragment targeted by dsRNA-HM05104 using gene synthesis (Integrated DNA Technologies). A DNA fragment encoding ND-42* was PCR-amplified to produce the pUbi-ND-42* transgene, that directs the constitutive expression of ND-42* under the control of the *Ubi-p63E* promoter, and the ato3’-ND-42*, that direct the expression of ND-42* under the control of an ato3’ enhancer (a 1828 nt DNA fragment located 2824 nt 3’ to the *ato* stop codon (Couturier et al., 2024)). The two plasmids above were obtained by Gibson assembly.

All DNA constructs produced by PCR and/or Gibson assembly were verified par sequencing. Primers and cloning details are available upon request.

### smiFISH

smiFISH probes were prepared at previously described (Tsanov et al., 2016). In short, FLAP-X oligonucleotide were annealed with the smiFISH probe set mix in Tris-HCl 50 mM pH = 7.5, NaCl 100 mM, MgCl2 10 mM using a thermocycler (85 °C, 3 min; 65 °C, 3 min; 25 °C 3 min). smi FISH probes were designed using Stellaris Inc. probe designer. The sequence of each probe set is available upon request. FISH was performed as described in (Trcek et al., 2017). Briefly, brain complexes were dissected from third instar larvae in phosphate-buffered saline (PBS) and fixed in 4% paraformaldehyde (in PBS) for 20 mins at room temperature (RT). After, samples were permeabilized using 0.5% Triton-X in PBS and transferred into 4M Urea in 2xSSC. Then the samples were hybridized at 37°C overnight in SSC 2×, Urea 4 M, Dextrane 10%, Vanydyl complex 10 mM, 0.15 mg/ml salmon sperm DNA, 100 μM smiFISH probe and anti-GFP (1:1000) and rinsed with 4M Urea in 2xSSC and after with 2xSSC. The secondary antibody was incubated 2hr in PBS with 0.1% Triton-X in (PBT) at RT.

### Immunostaining

Dissected brain complexes were fixed in 4% paraformaldehyde (in PBS) for 20 mins, washed in PBT, incubated 1-2 hr at RT in PBT with first antibodies, washed 3x 10 mins in PBT, incubated 1-2 hr at RT with secondary antibodies, washed with PBT and mounted in mounting medium (90 % glycerol in PBS, 0.1 % n-propyl gallate, Sigma P3130). The following primary antibodies were used: rat anti-Ecad (DSHB, 1:100); goat anti-GFP (Abcam, 1:1000); rabbit anti-dsRed ( 1:1000). The following secondary antibodies were used: Cy3 anti-rat (1:1000); Cy2 anti-goat (1:1000); Cy5 anti-rat (1:1000). Secondary antibodies were from Jackson Laboratories. After incubation, samples were briefly washed with PBT, followed by three 10 min washes in 0.1% PBT. Samples were calibrated in 50 % Glycerol/PBS, before eye-antennal imaginal discs were dissected and mounted in mounting media (10 % PBS 10X, 90 % glycerol (ACS grade 99-100% purity), 0.1 % n-propyl gallate (20% (w /v), Sigma P3130).

### *Ex vivo* culturing of eye discs

Third instar larvae were briefly washed in water, rinsed in PBS 1x and dissected in Grace’s medium (Sigma G9771) at pH 6.7 supplemented with 5% Fetal Bovine Serum (FBS), Penicillin/Streptomycin 0.5% (Sigma P4333) and 20 nM 20-Hydroxyecdysone (Sigma H5142). Eye-antenna imaginal discs were dissected and transferred into a magnetic imaging chamber (Chamlide CM-B25-1, LCI) with a drop of culture medium. Discs were then embedded using a fibrinogen-thrombin mix (Sigma 11424246 and 11407522), then cultured 20 mins at 25°C in supplemented Grace’s medium before imaging at 23-25°C. Antimycin A (10µM in 99% EtOH) and bromopyruvic acid (3BP, 100µM in imaging medium) were added to the medium directly in the imaging chamber.

### **M**icroscopy

Images were acquired on a confocal Zeiss LSM780 microscope equipped with 20x (PL APO, N.A. 0.8) and 40x (PL APO, N.A. 1.32) objectives and a Nikon Ti2-E AX-NSPARC microscope with a 40x (PL APO, N.A. 1.25) objectives. The iATPsnFR sensor was excited using the 488 and 561 nm lasers and the emitted light was collected in the 490-534 and 579-641 nm intervals. The Peredox-Cherrynls sensor was excited using the 405 and 561 nm lasers and the emitted light was collected between 409-540nm and 597-659nm, respectively. SoNar was excited using 488 nm and 564 nm laser and light was collected around 530 nm. The signals of these metabolic sensors were measured in UPs and DCs over user-defined regions of interest on average z-projected images using ImageJ.

Scanning Electron Microscopy (SEM) was performed on dehydrated adult flies coated with gold palladium following standard procedures.

### NAD^+^/NADH-glo assay

Total cellular NADH and NAD^+^ were measured using NAD^+/^NADH-glo assay (Promega). Around 40 eye discs were dissected in PBS and lysed using 1% DTAB (Sigma, D8638) in base solution (0.2N NaOH). Lysate fractions corresponding to 5 and 10 eye discs were pipetted into a 96-well plate (Corning 96 Flat Bottom white Polystyrene). The remaining lysate was stored at −20°C and used for protein quantification in which a standard curve was included for each experiment (BCA assay, Pierce). The manufacturer’s instructions were followed thereafter. Briefly, 50 µl of the lysate was then transferred to second well containing 25 µl of 0.4 M HCl (for NAD+ measurement) or to an empty well (for NADH measurement). Both samples were heated at 60 °C for 15 min to deplete NAD+ or NADH. Samples were then left on the bench for 10 min to cool down to room temperature before adding neutralizing solutions (25 µl of 0.5 M Tris base to the NAD+ sample, and 50 µl of 1:1 0.4 M HCl:0.5 M Tris base to the NADH sample), followed by 100 µl of NADH-Glo detection reagent. Following incubation for 1hr at room temperature, the produced luminescence signal was measured on a microplate reader (Tecan INFINITE 200 PRO). Absolute concentrations of NADH and NAD+ were determined against standard curves using NADH (Sigma, N8129) and NAD^+^ (Sigma N7004). Measured concentrations were normalized by protein concentration values.

### Targeted LC–MS metabolomics analyses

For each sample, eye-antenna discs from 30 female larvae were rapidly dissected in PBS at 4°C and frozen using dry ice. Metabolites were extracted using 0.1 mL of a solution composed of 50% methanol, 30% acetonitrile (ACN) and 20% water. Samples were then vortexed for 5 min at 4 °C and centrifuged at 16,000 g for 15 min at 4 °C. The supernatants were collected and stored at −80 °C until analysis. LC/MS analyses were conducted on a QExactive Plus Orbitrap mass spectrometer equipped with an Ion Max source and a HESI II probe coupled to a Dionex UltiMate 3000 UPLC system (Thermo). The 5 µl samples were injected onto a ZIC-pHILIC column (150 mm × 2.1 mm; i.d. 5 µm) with a guard column (20 mm × 2.1 mm; i.d. 5 µm) (Millipore) for LC separation. Buffer A was 20 mM ammonium carbonate, 0.1% ammonium hydroxide (pH 9.2), and buffer B was ACN. The chromatographic gradient was run at a flow rate of 0.2 µl min−1 as follows: 0–20 min, linear gradient from 80% to 20% of buffer B; 20–20.5 min, linear gradient from 20% to 80% of buffer B; 20.5–28 min, 80% buffer B. The mass spectrometer was operated in full scan, polarity switching mode with the spray voltage set to 2.5 kV and the heated capillary held at 320 °C. The sheath gas flow was set to 20 units, the auxiliary gas flow to 5 units and the sweep gas flow to 0 units. The metabolites were detected across a mass range of 75–1000 m/z at a resolution of 35,000 (at 200 m/z) with the automatic gain control target at 106 and the maximum injection time at 250 ms. Lock masses were used to ensure mass accuracy below 5 ppm. Data were acquired with Thermo Xcalibur software (Thermo). The peak area of metabolites was determined using Thermo TraceFinder software (Thermo), identified by the exact mass of each singly charged ion and by the known retention time on the HPLC column. To perform protein normalization, the remaining protein pellet was diluted in lysis buffer (1% DTAB (Sigma, D8638) in base solution (0.2N NaOH)) and used for protein quantification (BCA assay, Pierce). Data was analyzed using the MetaboAnalyst v6.0 (Pang et al., 2024).

### Western blot analysis

Eye-antenna discs from 20 female larvae were lysed using NuPAGE LDS sample buffer (Invitrogen). Inhibitors of PARG (1 μM; PDD00017273, Sigma), and PARP activities (1 μM; Olaparib, LKT LABS) were added in the lysis buffer. Following denaturation (100 °C for 10 min), proteins were separated on Mini-PROTEAN TGX Stain-Free Gels, 8-16% (Bio-Rad) and transferred to nitrocellulose membranes (Bio-Rad) overnight at 4 °C. The blotted membranes were blocked with 10% Bovine Serum Albumine (w/v) in Tris Buffer Saline (with 0.1% Tween 20 (TBS-Tw, 1 h at room temperature). The top (>55 kDa) and bottom (<35kDa) parts of the membranes were incubated with rabbit anti-poly-ADPr (MABE1031, Millipore, 1:500, RRID: AB_2665467), while the central part (35-55kDa) was incubated with rabbit anti-fibrillarin (ab5821, Abcam, 1:1000) for 1hr at room temperature. After washes in TBS-Tw, blots were incubated with anti-rabbit StarBright 700 (Bio-Rad). Signals were detected using a Bio-Rad Chemi-Doc MP Imager and analysed with the Bio-Rad Image Lab software.

### scRNAseq

Brain complexes from Oregon-R and *mirr > ND-42^RNAi^* larvae were dissected in Dulbecco’s phosphate-buffered saline (DPBS; ThermoFisher) and then dissociated in 0.25% trypsin-EDTA solution to a single cell suspension at 37°C for 10 min. These cells were then washed by DPBS twice and passed through a 35μm filter before library preparation. Construction of 10x single-cell libraries and sequencing on the Illumina Hiseq platform were performed by Novogene. Raw data mapping was performed using standard Cell Ranger pipeline (v 2.2.0) to generate UMI count matrices. The NCBI GEO accession number for this dataset is GSE294725. We performed read alignment and annotation using the BDGP6 genome reference fastaq file and the BDGP6.91.gtf file.

The cluster comprising eye field cells was identified based on *eya* expression. Cells from this cluster were further clustered using the Seurat package (v5.1.0) (Hao et al., 2021; Satija et al., 2015) and peripodial cells, UPs and DCs were identified based on previously described markers (Bravo González-Blas et al., 2020; Yeung et al., 2022). UPs and DCs were further annotated as v and d cells based on the expression of *mirror* (*mirr*), *araucan* (*ara*) and/or *caupolican* (*caup*). Differentially expressed genes (DEGs) were defined using the FindMarkers function. The d *vs* v DEGs found in Oregon-R were removed from the list of DEGs found in *mirr>ND-42^RNAi^* eye discs. Differentially expressed transposable elements were processed separately. GO term analysis was performed using the EnrichR package (Chen et al., 2013; Kuleshov et al., 2016; Xie et al., 2021) using the “GO_Biological_Process_2018” database.

## Legend of supplementary Figures and Tables

**Figure S1:**
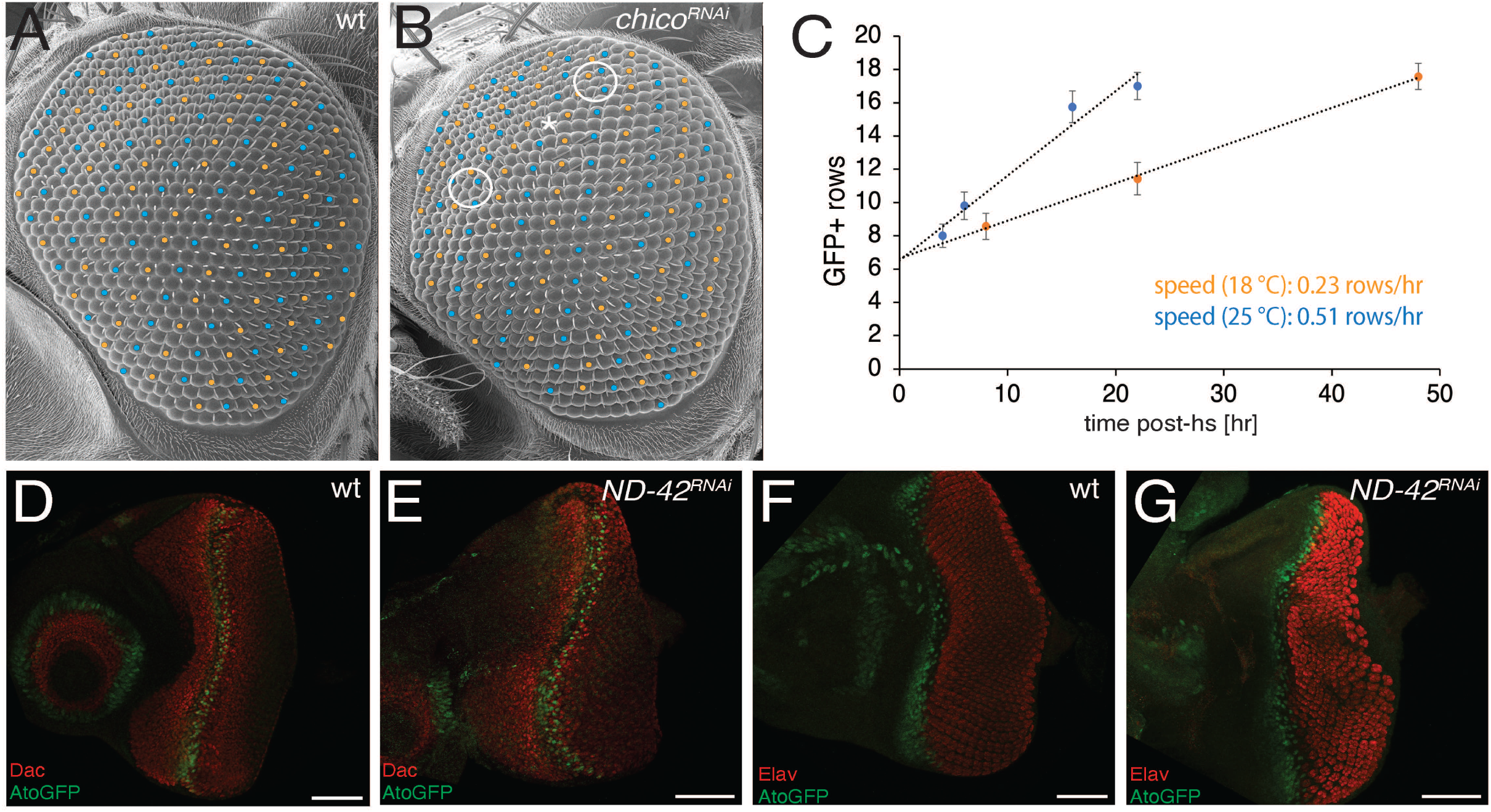
Patterning speed analysis. **A, B)** Scanning Electron Microscopy (SEM) images of adult fly eyes. A regular crystal-like array of ommatidia was observed in wt (A) whereas minor defects suggestive of interrupted rows were seen in the dorsal part of the eye in *mirr>*c*hico^RNAi^* flies (B; white circles indicate where non-consecutive rows appeared to merge; the star indicate a partial interruption). **C)** The speed of progression of the MF was measured in control *mirr>mIFP* discs from larvae grown at 18°C (orange) and 25°C (blue) using a recombination-based RFP-to-GFP switch. The MF moved twice slower at 18°C (0.23 rows/hr) than at 25°C (0.51 rows/hr). **D-G)** Dachsund (Dac, red in D,E) and Elav (red, in F,G) showed similar pattern of expression in control *mirr-Gal4 ato^GFP^* (D,F) and *mirr>ND-42^RNAi^ ato^GFP^* eye discs (E,G). AtoGPP (green) marked the position of the differentiation front. Scale bars, 50µm.

**Figure S2:**
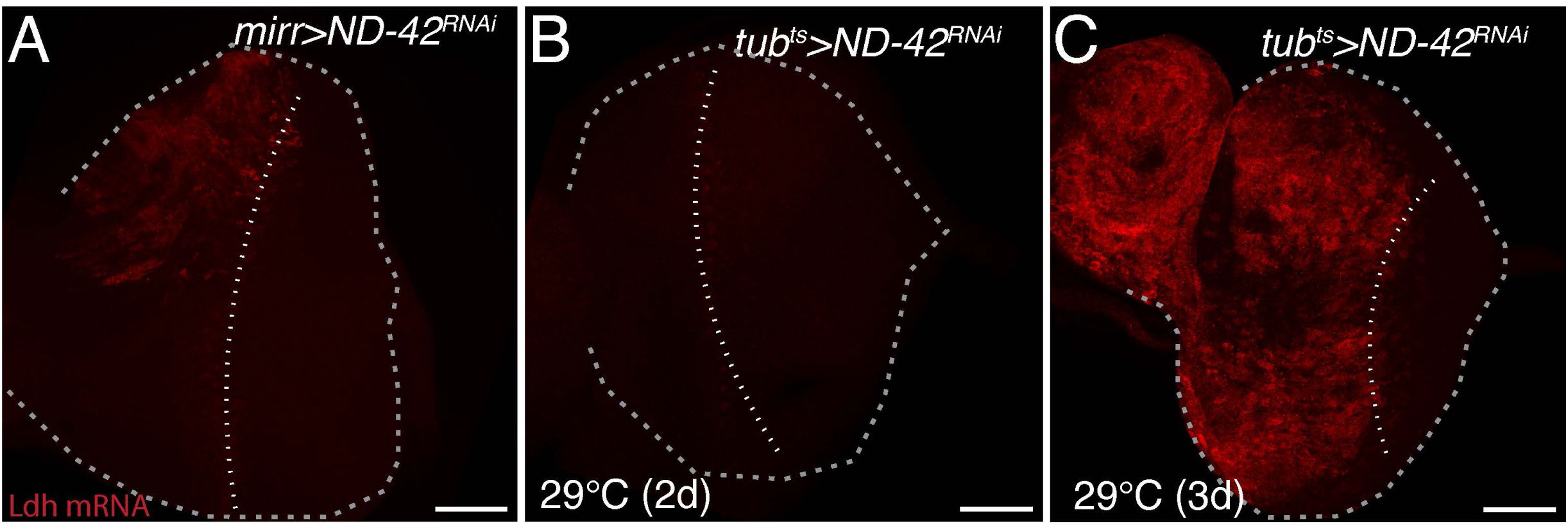
Conditional silencing of the *ND-42* gene. Conditional silencing of the *ND-42* gene using a *tub-Gal4 tub-Gal80^ts^*driver (*tub^ts^*) led to *Ldh* mRNA accumulation (red) in UPs, peripodial and antenna cells after 3 days (3d) at 29°C (C). This effect appeared to be stronger than the one observed using *mirr-Gal4* (A). The *Ldh* signal was very weak after 2 days at 29°C (B). The position of the MF was indicated by a white dotted line (disc outlines, grey dotted lines). Scale bars, 50µm.

**Figure S3:**
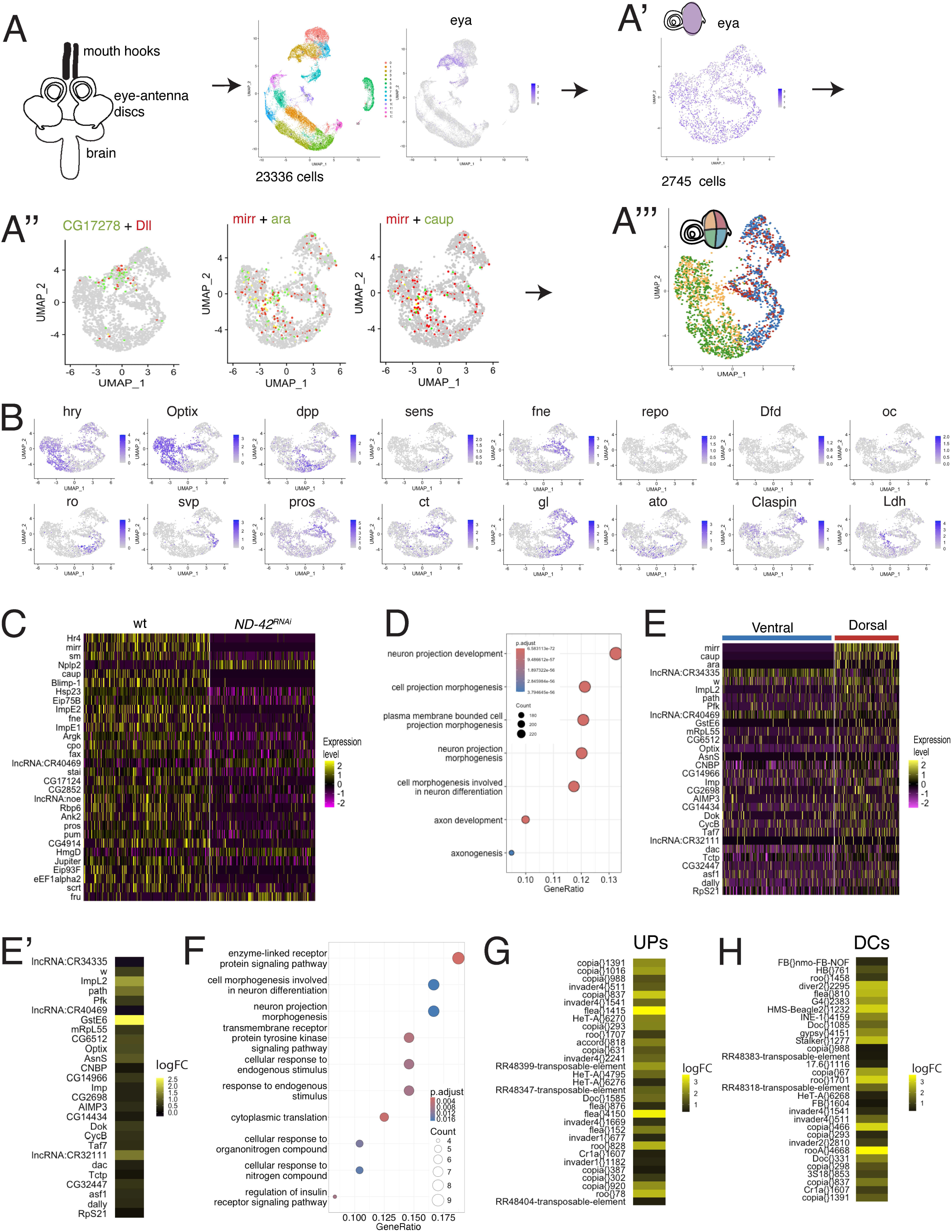
A scRNAseq analysis of the DC response to ETC inhibition. **A-B)** scRNAseq was performed on cells from dissected eye-brain complexes from control Oregon-R larvae (30,045 cells) and *mirr>ND-42^RNAi^* larvae (23,336 cells). Clustering analysis identified one cluster marked by *eya*+ cells corresponding to the eye primordium (A,A’). Cells from this cluster (2,745 and 2,112 cells for the *mirr>ND-42^RNAi^*and Oregon-R samples, respectively) were further clustered (A’’), leading to the identification of a peripodial cell cluster based on *DII* and *CG17278* gene expression (A’’). This cluster was removed in all further analysis. The remaining UPs (1,275 cells; color coded green and orange) and DCs (1,127 cells; color coded red and blue) were identified based on marker gene expression (A’’’,B). These eye cells were also categorized as *d* and *v* cells based on the expression of the *mirr, ara* or *caup* genes (triple-negative cells were identified as *v* cells; see schematic in A’’’ for the color code). **C)** Heatmap of the top 30 differentially expressed genes between wild-type *v* and *d* cells (wt, Oregon-R) and *mirr>ND-42^RNAi^* control *v* cells (gene expression level color-coded as indicated). Genes were ranked by adjusted p-values. A few ecdysone-regulated genes and differentiation markers appeared to be differentially expressed between these two populations of control cells. **D)** Gene Ontology (GO) term enrichment analysis for biological pathways for the top 1781 genes. **E,E’)** The top 30 differentially expressed genes between *ND-42^RNAi^ d* DCs (n=414) and control *v* DCs (n=713) DCs from *mirr>ND-42^RNAi^*discs were ranked by adjusted p-values and expression levels were color coded (E; see heatmap). Relative changes in averaged expression levels are shown as log2 Fold hange values (logFC; E’). The three genes used for clustering (*mirr*, *ara* and *caup*) were removed from the analysis shown in E’. **F)** GO term enrichment analysis for biological pathways for the top 60 genes with p-value <0.05 in DCs. **G,H)** The top 30 differentially expressed transposable elements (TE) in DCs (n=73 TEs) and UPs (n=53 TEs) were ranked by p-values and expression levels (logFC values) were color coded as indicated.

**Figure S4:**
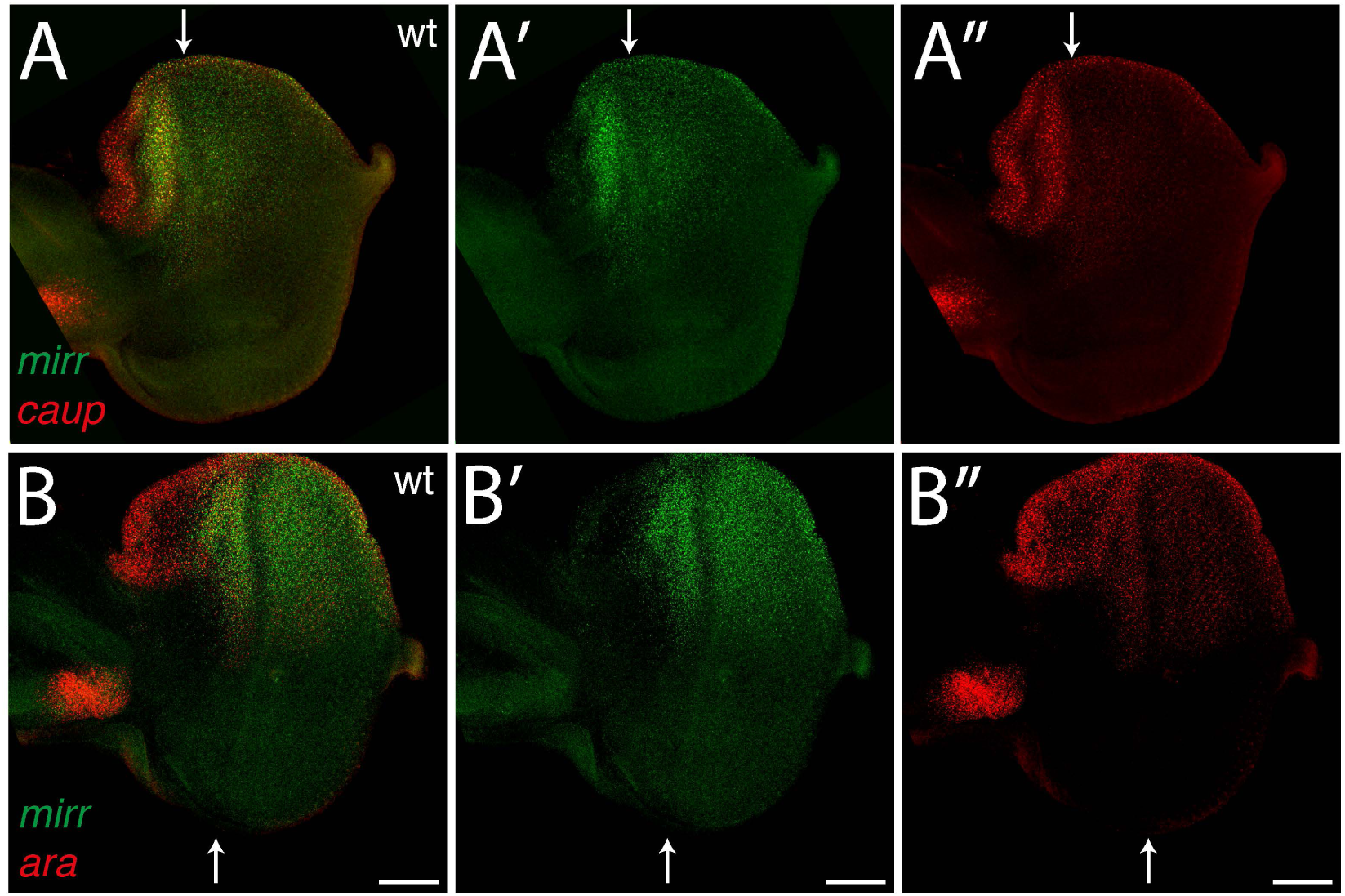
Co-expression of the *ara*, *caup* and *mirr* genes. Expression pattern of the *ara* (red in A,A’’), *caup* (red in B,B’’) and *mirr* genes (green in A-B’) showing that all three genes were largely co-expressed in *d* eye field and peripodial cells (MF position indicated by white arrows).

**Figure S5:**
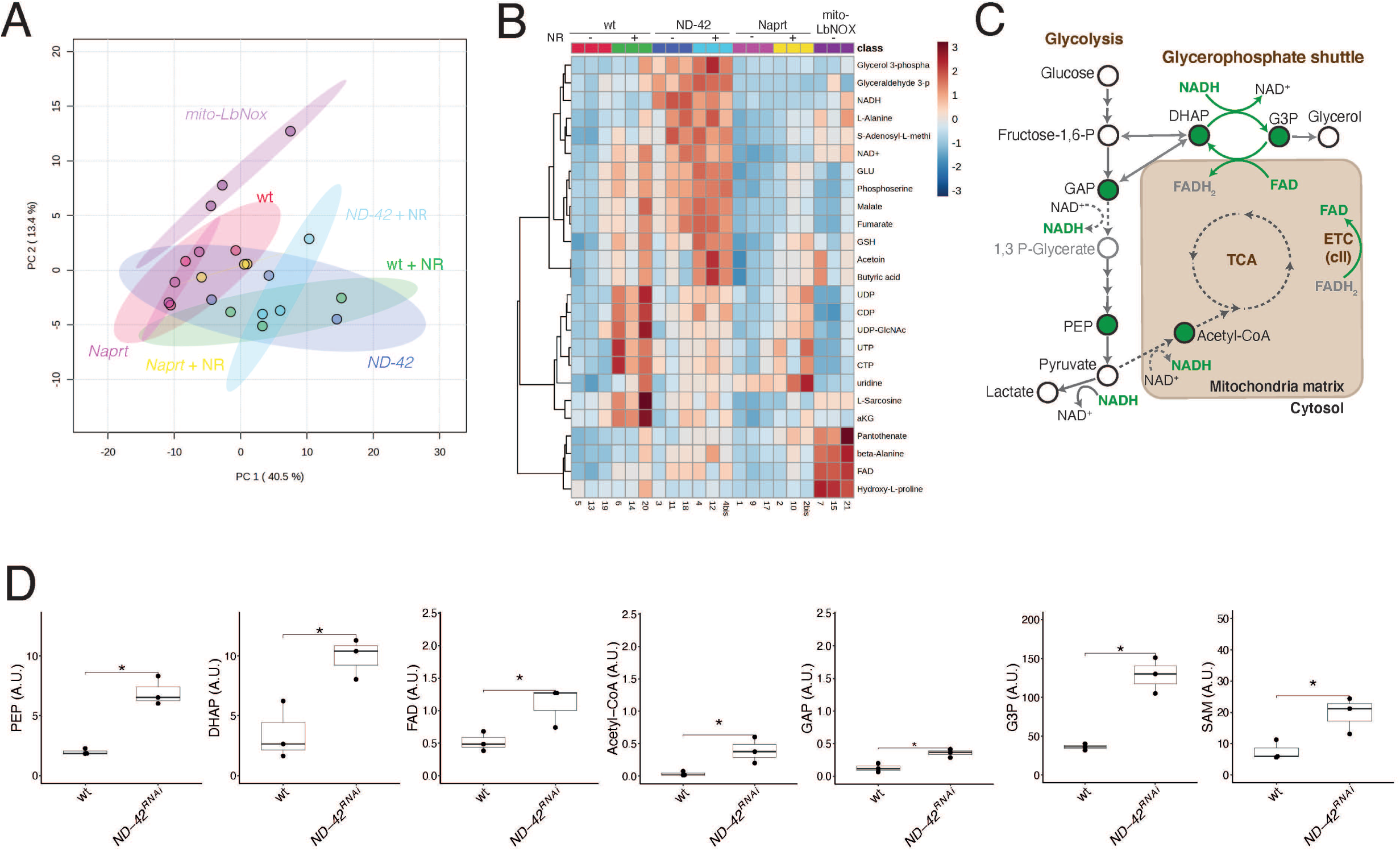
A metabolomic analysis of cI inhibition and reduced NAD^+^ recycling by LC/MS. **A)** PCA analysis of the metabolome (149 metabolites) of whole cell extracts prepared from *tub^ts^*>*+, tub^ts^>ND-42^RNAi^, tub^ts^*>*Naprt^RNAi^*and *tub^ts^*>*mito-LbNOX* eye discs. These discs were dissected from larvae grown in fly food supplemented or not with NR (as indicated). Clustering analysis indicated that the *tub^ts^*>*Naprt^RNAi^*metabolome was distinct from those of all other samples. Also, the *tub^ts^>ND-42^RNAi^* metabolome differed from the *tub^ts^*>*+* control metabolome in the absence of NR supplementation. In the presence of NR, the effect of *ND-42^RNAi^* appeared to be more restricted. **B)** Clustering analysis based on the top 25 differentially detected metabolites (abundance plotted as a heatmap). Differences in NAD, glycolysis anf TCA cycle metabolism were detected in the *tub^ts^>ND-42^RNAi^* metabolome relative to the *tub^ts^*>*+* control metabolome. **C)** Proposed changes in energy metabolism pathways based on the LC/MS results. Metabolites that showed a statistically significant increase upon *ND-42^RNAi^* relative to wild-type condition appear in green (see Table S2 for FC values). Enzymatic reactions proposed to be up-regulated are shown as green arrows; those proposed to be down-regulated appear as dotted arrows. Limiting NAD^+^ levels is proposed to result in substrate accumulation in reactions coupled to NAD^+^ reduction, e.g. GAP. Increased DHAP and GAP levels are interpreted to suggest increased Glycerol Phosphate shuttle activity, promoting NAD^+^ regeneration in the cytosol and reduction of Flavin Adenine Dinucleotide (FAD) in mitochondria (FADH_2_ could then serve to fuel the ETC via the cII). High levels of Acetyl-CoA is proposed to accumulate due to defective TCA activity. **D)** Plots showing the LC/MS results for the PhosphoEnolPyruvic acid (PEP), DHAP, FAD, Acetyl-CoA, GAP,G3P and SAM. Wilcoxon test: *< 0.05.

**Figure S6:**
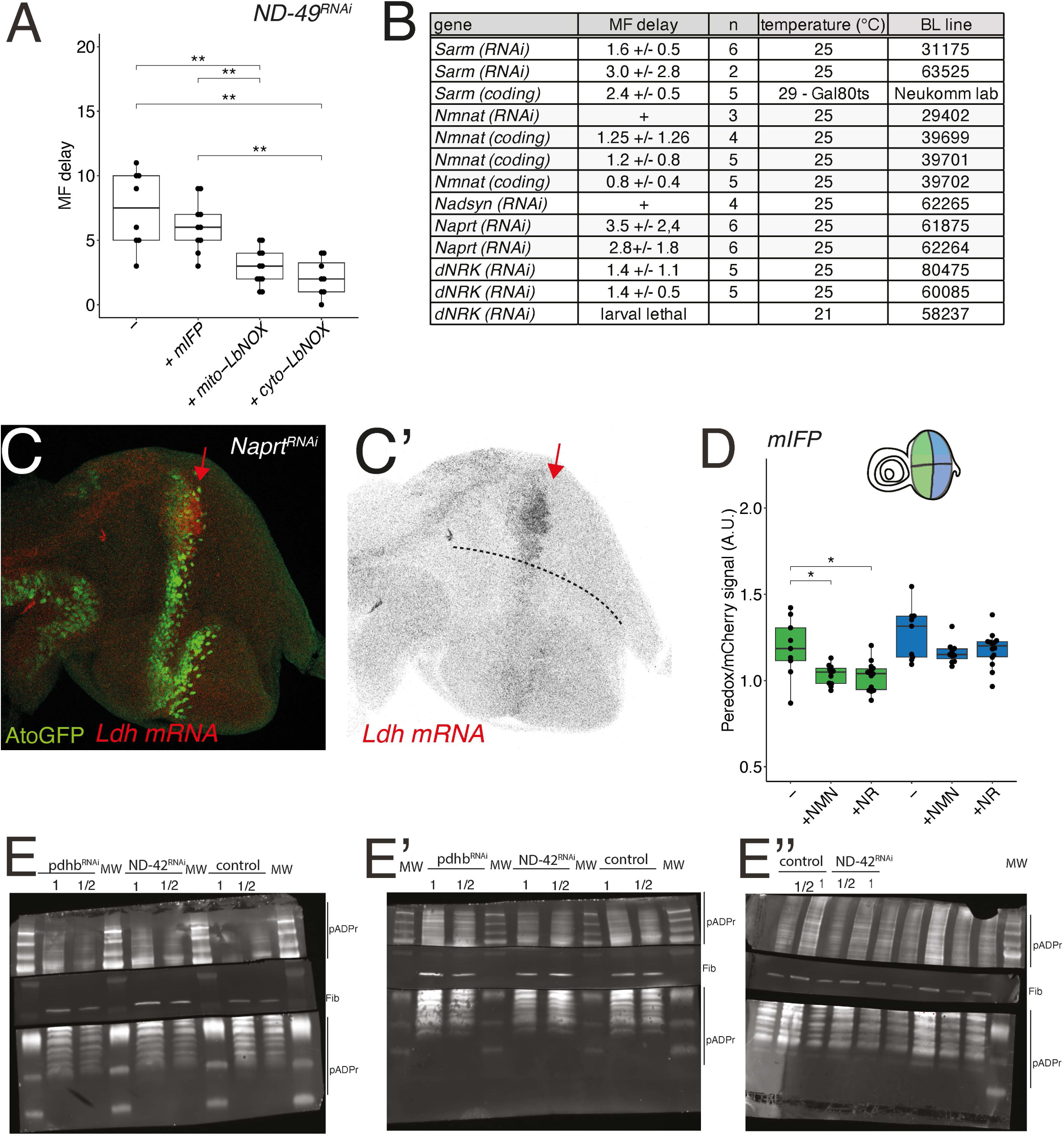
NAD^+^, cellular redox and speed of progression of the MF. **A)** MF delay analysis showing that the expression of cytosolic or mitochondrial LbNOX was sufficient to reduce the delay resulting from the silencing of the *ND-49* gene: *mirr>ND-49^RNAi^*: 7.0 +/−3.4 (n=5); *mirr>ND-49^RNAi^ + mIFP*: 6.1 +/−2.0 (n=10); *mirr>ND-49^RNAi^ + mito-LbNOX*: 2.9 +/−1.4 (n=11); *mirr>ND-49^RNAi^ + cyto-LbNOX*: 2.1 +/−1.5 (n=8). **B)** MF delay analysis of NAD metabolism genes (see Fig 7C). ‘+’ indicated that the number of ommatidial rows were not counted as discs looked clearly similar to wild-type. **C,C’)** Loss of Naprt activity in *mirr>Naprt^RNAi^ ato^GFP^*discs led to a compensatory change in *Ldh* gene expression (red; AtoGFP, green). The position of the MF is indicated by a red arrow. The position of the *dv* boundary is indicated by a dotted line. **D)** Analysis of the Peredox normalized intensity ratio in *mirr>mIFP* eye discs indicated that addition of NMN and NR lowered the cellular redox in UPs, but not DCs. Wilcoxon test: *< 0.05. **E-E’’)** Western Blot analysis of control, *ND-42^RNAi^* and *pdhb^RNAi^* eye discs. Fibrillarin (Fib; blots at center) was used to normalize the poly-ADPr signal (blots at top and bottom). Each sample was loaded twice (with a 2-fold difference in volume; MW, Molecular Weight).

**Table S1: Results of the perturbation screen**

MF delay values (mean and s.d.; n represents the number of discs; NA: not scorable; +: no phenotype and not scored), temperature condition, lethality (L, larval; P, pupal) and stock numbers are indicated. An arbitrary threshold value (>2; highlighted in blue) was set to identify relevant perturbations. The 166 genes studied here appear in broad functional categories. Severe perturbations resulting in disorganized ommatidia, tissue loss and/or lethality prevented us from studying MF progression (NA). Some of the corresponding genes were re-tested at lower temperature and/or in combination with Gal80ts to reduce the strength of the perturbation. For instance, while a strong loss of Complex IV (*cype* gene) or ATP synthase (*blw* gene) activity resulted in lethality (at 25°C), a weaker loss of function produced a delayed MF phenotype which was reported in the table. Beyond energy metabolism, perturbations affecting various aspects of cellular metabolism were identified, including Rbf, a transcription factor regulating cellular metabolism; Zw, the first and rate limiting enzyme of the pentose phosphate pathway that uses sugar to produce nucleotides; Dpck, a kinase involved in CoA biosynthesis; and Gnmt, a glycine N-methyltransferase, that regulates the biosynthesis of the S-Adenosyl Methionine, a methyl donor.

**Table S2: LC/MS results**

For each metabolite measured by LC/MS, the raw value (sheet #1, raw values), as well as the FC and p-values of the *ND-42^RNAi^ vs* wild-type comparison (sheet #2), are provided. Each genotype was studied in triplicate with the exception of *Naprt^RNAi^*+NR and *ND-42^RNAi^*+NR (in both cases, one samples was analyzed twice by LC/MS, as noted with ‘bis’). Raw values were normalized by protein concentration measured by BCA (line 3, grey).

**Table S3: Differentially expressed genes in *mir>ND-42^RNAi^* discs**

Lists of genes that were differentially expressed between *d* and *v* cells, for both UPs and DCs, ranked by adjusted p-values. The log2FC and the percentage of cells for which the corresponding gene was found to be expressed in the *d* (ptc.1) and *v* (ptc.2) categories are also shown.

**Table S4: Differentially expressed Transposable Elements in *mir>ND-42^RNAi^* discs**

TEs (FlyBase ID and name) that were differentially expressed between *d* and *v* cells, for both UPs and DCs, were ranked by p-values. The log2FC and the percentage of cells for which the corresponding gene was found to be expressed in the *d* (ptc.1) and *v* (ptc.2) categories are also shown.

